# SARS-CoV-2 B Epitope-Guided Neoantigen NanoVaccines Enhance Tumor-Specific CD4/CD8 T Cell Immunity Through B Cell Antigen Presentation

**DOI:** 10.1101/2024.07.18.604163

**Authors:** Chengyi Li, Shuai Mao, Zera Montemayor, Mohamed Dit Mady Traore, Alejandra Duran, Fang Ke, Mahamadou Djibo, Hanning Wen, Wei Gao, Duxin Sun

**Affiliations:** Department of Pharmaceutical Sciences, College of Pharmacy, University of Michigan, Ann Arbor, Michigan 48109, United States

## Abstract

Current neoantigen cancer vaccines activate T cell immunity through dendritic cell /macrophage-mediated antigen presentation. It is unclear whether incorporating B cell-mediated antigen presentation into current neoantigen vaccines could enhance CD4/CD8 T cell immunity to improve their anticancer efficacy. We developed a SARS-CoV-2 B cell epitope-guided neoantigen peptide/mRNA cancer nanovaccines (B_SARS_T_NeoAg_Vax) to improve anticancer efficacy by enhancing tumor-specific CD4/CD8 T cell antitumor immunity through B cell-mediated antigen presentation. B_SARS_T_NeoAg_Vax crosslinked with B cell receptor, promoted SARS-CoV-2 B cell-mediated antigen presentation to tumor-specific CD4 T cells, increased tumor-specific follicular/non-follicular CD4 T cells, and enhanced B cell-dependent tumor-specific CD8 T cell immunity. B_SARS_T_NeoAg_Vax achieved superior efficacy in melanoma, pancreatic, and breast cancer models compared to the current neoantigen vaccines. Our study provides a universal platform, SARS-CoV-2 B epitope-guided neoantigen nanovaccines, to improve anticancer efficacy against various cancer types by enhancing CD4/CD8 T cell antitumor immunity through viral-specific B cell-mediated antigen presentation.

## INTRODUCTION

Neoantigen cancer vaccines, which use CD4/CD8 T cell neoantigens from individual cancer patients to activate CD4/CD8 T cell antitumor immunity, have shown promising anticancer efficacy in patients with melanoma and pancreatic cancer.^1–4^ It remains to be seen if these neoantigen cancer vaccines are effective in patients with other types of cancers,^1,5–18^ and thus there is critical needs to develop strategies to further enhance their efficacy in different types of cancers.

Current neoantigen peptide or mRNA cancer vaccines mainly utilize dendritic cell (DC)/macrophage-mediated antigen presentation to activate CD4/CD8 T cell antitumor immunity.^1,5–23^ Despite B cells being professional APCs,^24–26^ these vaccines do not utilize B cell-mediated antigen presentation due to the longstanding debate for the role of B cell immunity in cancer.^27–35^ However, recent studies suggest that B cell-mediated antigen presentation by tumor-infiltrating B cells (TIL-Bs) may promote tumor-specific CD4/CD8 T cell antitumor immunity.^36–44^ Clinical studies show that higher B cell infiltration and activation in tumors correlates with better patient survival and improved response to anti-PD-1/PD-L1 immunotherapy across various cancer types.^38,45–49^ These evidences raise the question of whether incorporating B cell-mediated antigen presentation into current neoantigen vaccines could enhance CD4 and CD8 T cell immunity to improve their anticancer efficacy.

Utilizing B cell-mediated antigen presentation in cancer vaccines may offer unique advantages for activating T cell responses, distinct from DC/macrophage-mediated antigen presentation. First, while dendritic cells and macrophages capture antigens through non-specific binding, B cells use their surface immunoglobulin molecules (B cell receptors, BCRs) to specifically and efficiently bind and uptake cognate antigens with B cell epitopes. The binding of specific B cell epitopes to BCRs leads to BCR crosslinking, which subsequently enhances peptide antigen uptake, processing, and presentation via peptide-loaded MHC-II complexes to activate CD4 T cells.^24,25^ This unique process makes B cells more effective than dendritic cells and macrophages in presenting low-abundance antigens to T cells.^26^ This may provide advantages since antigen-specific B cells can effectively capture large amounts of antigen through high-affinity binding and provide stronger TCR stimulation.^50,51^ Additionally, B cells are the only professional APCs capable of clonal expansion, enabling them to direct the immune response towards specific antigens.^24,25,52^ The expanded number of B cells can in turn uptake, process, and present antigens to activate antigen specific CD4 T cells. ^50,51^

However, it is challenging to incorporate B cell-mediated antigen presentation into current neoantigen vaccines. To achieve this, the vaccine must include two components: (1) B Cell epitopes applicable to various cancers need to be included in the vaccines: However, current neoantigen cancer vaccines only include CD4/CD8 T cell epitopes derived from patients’ tumors, with no reported B cell neoantigens identified.^53,54^ We and others have shown that B cell epitopes from tumor-associated antigens (TAA), such as HER-2, VEGF, and EGFR, can be used to enhance B/CD4 T cell crosstalk in cancer vaccines.^55,56^ However, B cell epitopes from tumor-associated antigens (TAA) are only suitable for a small portion of patients with high antigen expression and are not adaptable to personalized neoantigen vaccines for various cancers.^57^ (2) Specialized delivery system needs to be used: A nano-sized carrier featuring multivalent B cell peptide epitopes on its surface may enhance BCR crosslink, antigen uptake, and the B cell-mediated antigen presentation. Nano-sized carrier may enhance lymph node delivery, allowing B cell epitopes to encounter B cells.^58^ The multivalent B cell epitopes increase the affinity of antigen binding to BCR receptor and facilitate the crosslinking of B cell receptors (BCR) resulting in efficient uptake and presentation of tumor antigens via the MHC-II complex to CD4 T cells.^25,59–62^

In this study, we developed a SARS-CoV-2 B cell epitope-guided neoantigen peptide or mRNA cancer nanovaccine (B_SARS_T_NeoAg_Vax or B_SARS_T_mRNA_Vax) to promote tumor-specific CD4/CD8 antitumor immunity through B cell-mediated antigen presentation. We aim to test the hypothesis that B_SARS_T_NeoAg_Vax or B_SARS_T_mRNA_Vax could enhance tumor specific CD4/CD8 T cell immunity through SARS-CoV-2 specific B cell-mediated antigen presentation to tumor specific CD4 T cells, thereby improving anticancer efficacy in different types of cancers. We first generated a SARS-CoV-2 B cell epitope-guided neoantigen peptide nanovaccine (B_SARS_T_NeoAg_Vax) by conjugating both SARS-CoV-2 B cell epitopes and tumor T cell neoantigens on the surface of the nanocarrier.^55^ B_SARS_T_NeoAg_Vax enhanced anticancer efficacy compared to the neoantigen vaccine with only T cell epitopes in four different tumor models. B_SARS_T_NeoAg_Vax activated tumor-specific follicular and non-follicular CD4 T cells, and enhanced B cell-dependent tumor-specific CD8 T cell immunity. Furthermore, we also designed a SARS-CoV-2 B epitope-guided neoantigen mRNA cancer vaccines, which used a lipid nanoparticle (LNP) with multivalent SARS-CoV-2 B peptide epitopes on its surface, to encapsulate T cell mRNA neoantigen (P53R172H and KRAS G12D, B_SARS_T_KPC-mRNA_Vax). B_SARS_T_KPC-mRNA_Vax also enhanced anticancer CD4 and CD8 T cell immunity and showed better anticancer efficacy in pancreatic cancer model compared with LNP encapsulated with T cell mRNA neoantigen without B cell antigen presentation function. Altogether, our study demonstrates that SARS-CoV-2 B cell epitope-guided neoantigen cancer nanovaccines improved anticancer efficacy by enhancing tumor-specific CD4/CD8 T cell immunity through viral-specific B cell-mediated antigen presentation. Our study also provides a universal nanovaccine platform with viral B cell epitopes in neoantigen cancer vaccine design, potentially enhancing anticancer efficacy against different types of cancers.

### Prepare SARS-CoV-2 B epitope-guided neoantigen peptide NanoVaccines to promote BCR crosslink and B cell antigen presentation

We generated four types of NanoVaccines (**Fig. 1 A**).by conjugating viral antigen-cluster mimicry nanoparticles (ACN) with SARS-CoV-2 B epitope (SP14P5: CDDDTESNKKFLPFQQFGRDIA, SP21P2: CDDDPSKPSKRSFIEDLLFNKV) and tumor antigen, which include HER2^+^ breast cancer (tumor HER2, MHC-I-restricted peptide KIFGSLAFL and MHC-II-restricted peptide PESFDGDPA), B16-OVA (OVA antigen, MHC-I-restricted OVA_257-264_ peptide and MHC-II-restricted OVA_323-339_), melanoma (neoantigen, MHC-I-restricted M27: CDDDREGVELCPGNKYEMRRHGTTHSLVIHD^63,64^, and MHC-II-restricted M30: CDDDPSKPSFQEFVDWENVSPELNSTDQPFL), pancreatic cancer (neoantigen, Kras G12D: CDDDMTEYKLVVVGADGVGKSALTIQL, and Tp53 R172H: CDDDYKKSQHMTEVVRHCPHHERCSDGDG)^63,64^.

**Fig. 1.**
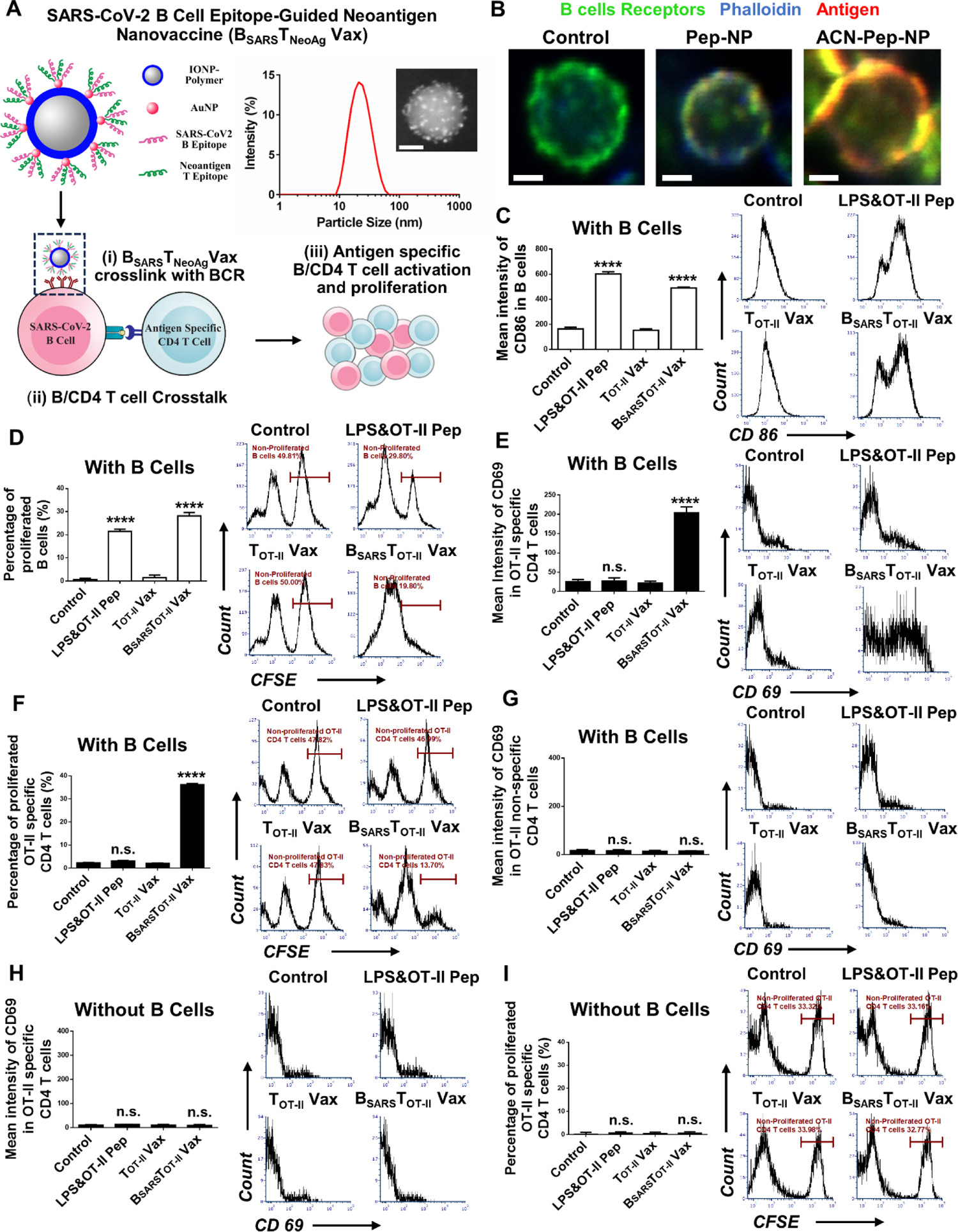
SARS-CoV-2 B epitope-guided T cell NanoVaccines promote BCR crosslink and B cell presentation to activate antigen specific CD 4 T cells. (A) Schematic illustration of the SARS-CoV-2 B Cell Epitope-Guided Neoantigen Nanovaccines (B_SARS_T_NeoAg_Vax). Size distribution of B_SARS_T_NeoAg_ Vax with dynamic light scattering (DLS) analysis and scanning transmission electron microscopy (STEM) high-angle annular dark-field (HAADF) image; Schematic illustration of the B_SARS_T_NeoAg_ Vax to promote BCR crosslink (i) and B cell antigen presentation mediated B/CD4 T cell crosstalk (ii), leading to the activating of antigen specific B cell and CD 4 T cell (iii). (B) BCR crosslink by B_SARS_T_NeoAg_Vax. Confocal image of Cy3 and hapten labeled ACN-Pep-NP (red) binding/crosslinking (yellow) with B cell receptor (antibody staining, green) in hapten-specific B cells from QM mice splenocytes, compared with Cy3 and hapten labeled peptide antigen (Pep-NP). Blue, phalloidin stain of actin filaments; green, B cell receptor staining using Alexa Fluor 488-AffiniPure Fab Fragment Goat Anti-Mouse IgM (µ Chain Specific) antibody; red: Cy3-labeled peptide-hapten. The scale bar is 2.5 µm. (C-I) B and CD 4 T cell crosstalk. Flow cytometry quantification and representative analysis (24 h) of B cell activation (C) from spike protein immunized mice by measuring the geometric mean intensity of CD86 marker. Flow cytometry quantification and representative analysis (96 h) of B cell proliferation from spike protein immunized mice, calculated by the decreased percentage of CFSE^+^ B cells before and after incubation (D). Flow cytometry quantification and representative analysis (24 h) of OT-II specific CD4 T cell activation with B cell incubation by measuring the geometric mean intensity of CD 69 marker (E). Flow cytometry quantification and representative analysis (96 h) of OT-II specific CD4 T cell proliferation with B cell incubation, calculated by the decreased percentage of CFSE^+^ OT-II specific CD4 T cells before and after incubation (F). Flow cytometry quantification and representative analysis (24 h) of OT-II non-specific CD4 T cell activation with B cell incubation by measuring the geometric mean intensity of CD69 marker (G). (H) Flow cytometry quantification and representative analysis (24 h) of OT-II specific CD4 T cell activation without B cell incubation by measuring the geometric mean intensity of CD69 marker. (I) Flow cytometry quantification and representative analysis (96 h) of OT-II specific CD4 T cell proliferation without B cell incubation, calculated by the decreased percentage of CFSE^+^ OT-II specific CD4 T cells before and after incubation. Data for quantification are shown as mean ± SD, n = 3. For **Figure 1C-1I**, statistical comparisons were conducted among B_SARS_T_OT-II_ Vax group and LPS&OT-II Pep group with other groups. Statistical comparisons are based on one-way ANOVA, followed by post hoc Tukey’s pairwise comparisons or by Student’s unpaired T-test. The asterisks denote statistical significance at the level of * p < 0.05, ** p < 0.01, *** p < 0.001, **** p < 0.0001. ANOVA, analysis of variance; SD, standard deviation; n.s., no statistical significance.

We generated viral antigen-cluster mimicry nanoparticles (ACN) with proven ability to crosslink with BCR and promote B cell antigen presentation for B and CD 4 T cell crosstalk (**Fig. 1 A**).^65^ ACN consists of an iron nanoparticle core (IONP, 15 nm), with ultra-small gold nanoparticles (AuNPs, 2 nm) attached to the surface. SARS-CoV-2 B peptide epitopes^66^, and T cell epitopes with cysteine group were linked to the surface AuNPs of the ACN through a thiol-Au reaction to achieve high density multivalent antigen cluster structure for BCR crosslink (**Fig. 1A**).^55,67–69^ After conjugation with antigens, the size of the B_SARS_T_NeoAg_Vax was 44 ± 2 nm by dynamic light scattering (DLS) with a polydispersity index (PDI) of 0.087 ± 0.03 (**Fig. 1A**).

To promote B cell antigen presentation, a vaccine must facilitate B cell epitope binding and crosslinking BCR, which subsequently enhances peptide antigen uptake, processing, and presentation via peptide-loaded MHCII complexes to activate CD4 T cells (**Fig. 1A**).^24,25^ To confirm the ability of B_SARS_T-Vax to facilitate the binding and crosslinking of B cell receptor (BCR), we conjugated fluorescent Cy3 and hapten labeled antigen to ACN nanoparticle (ACN-Pep-NP) and incubated with NP-specific B cells (NP will specially bind to hapten-antigen) from QM transgenic mice in comparison with free peptide-hapten (Pep-NP). ACN-Pep-NP showed significantly increased BCR crosslinking efficiency compared to Pep-NP using co-localization staining of ACN-Pep-NP (red) and BCR (green) (**Figure 1B and S1**).

To investigate whether B_SARS_T-Vax could promote SARS-B cell and CD 4 T cell crosstalk, we conjugated SARS-CoV-2 B cell epitope and OT-II specific CD4 T cell epitope (chicken ovalbumin_323-339_) with ACN (B_SARS_T_OT-II_Vax), incubated with B cells from lymph nodes and spleen of SARS-CoV-2 spike protein immunized mice with enriched SARS-CoV-2 B epitope-specific B cells, as well as OT-II specific CD4 T cells from the spleen of OT-II transgenic mice for 24 h or 96 h (**Figure 1C-1I, S2-S4**). LPS mixed with OT-II CD4 T cell epitope (LPS&OT-II Pep) and OT-II CD4 T epitope conjugated with ACN nanoparticle (T_OT-II_Vax) were used as comparison. We firstly monitored the activation and proliferation of B cells from spike protein immunized mice (B cells labeled with CFSE tracker). B_SARS_T_OT-II_Vax induced 3.2-fold (as monitored by CD86 intensity) and 15.6-fold (as monitored by CD69 intensity) higher activation and 33.5-fold higher proliferation (as monitored by CFSE intensity) of B cells than T_OT-II_ Vax (**Figure 1C, 1D and S2**).

To investigate whether B_SARS_T_OT-II_Vax activated B cells could process and present OT-II CD4 T cell epitope to activate OT-II specific CD4 T cells, we measured the activation and proliferation of OT-II specific CD4 T cells (labeled with CFSE tracker) by flow cytometry. B_SARS_T_OT-II_Vax stimulated 9.4-fold (as monitored by CD69 intensity) and 2.2-fold (as monitored by CD25 intensity) higher activation and 181.6-fold higher proliferation (as monitored by CFSE intensity) of OT-II specific CD4 T than T_OT-II_Vax (**Figure 1E, 1F and S3A**). As controls, no statistical differences of activation and proliferation were observed in antigen nonspecific CD4 T cells (not OT-II CD4 T cell epitope specific) between B_SARS_T_OT-II_Vax and T_OT-II_Vax (**Figure 1G and S3B**), which suggest the antigen specific activation of CD4 T cells was mediated by B_SARS_T_OT-II_ Vax activated B cells.

To further confirm the activation of OT-II specific CD4 T cell by B_SARS_T_OT-II_ Vax is B cell dependent, we depleted B cells from splenocytes and lymph node cell suspension and mixed them with OT-II specific CD4 T cells. Results showed that depletion of B cells severely impaired the activation and proliferation of OT-II specific CD4 T cells (**Figure 1H, 1I, and S4**). To further confirm that B_SARS_T_OT-II_Vax induced OT-II specific CD4 T cell is dependent on B cell antigen presentation, we mixed B cells from lymph node and spleen of spike protein immunized mice, OT-II specific CD4 T cells with αMHC-II, αCD40L or αICOS antibodies to block B and CD 4 T cell crosstalk. The blocking of B/CD 4 T cell crosstalk significantly eliminated the activation and proliferation of OT-II specific CD4 T cells by B_SARS_T_OT-II_Vax (**Figure S2-S4**). These results suggest that B_SARS_T_OT-II_Vax promotes the B cell antigen presentation to activate OT-II specific CD4 T cells.

### SARS-CoV-2 B epitope-guided cancer NanoVaccines induced SARS-CoV-2 specific B cell and tumor specific CD4 T cell activation in vivo

In order to promote B cell antigen presentation, the nanovaccine needs to efficiently deliver the B cell epitopes to lymph nodes to encounter B cells. We first confirmed that ACN with SARS-CoV-2 B peptide epitopes linked on the surface deeply penetrated inside the lymph node (**Figure 2A**), increasing the probability of antigens encountering B cells.

**Fig. 2.**
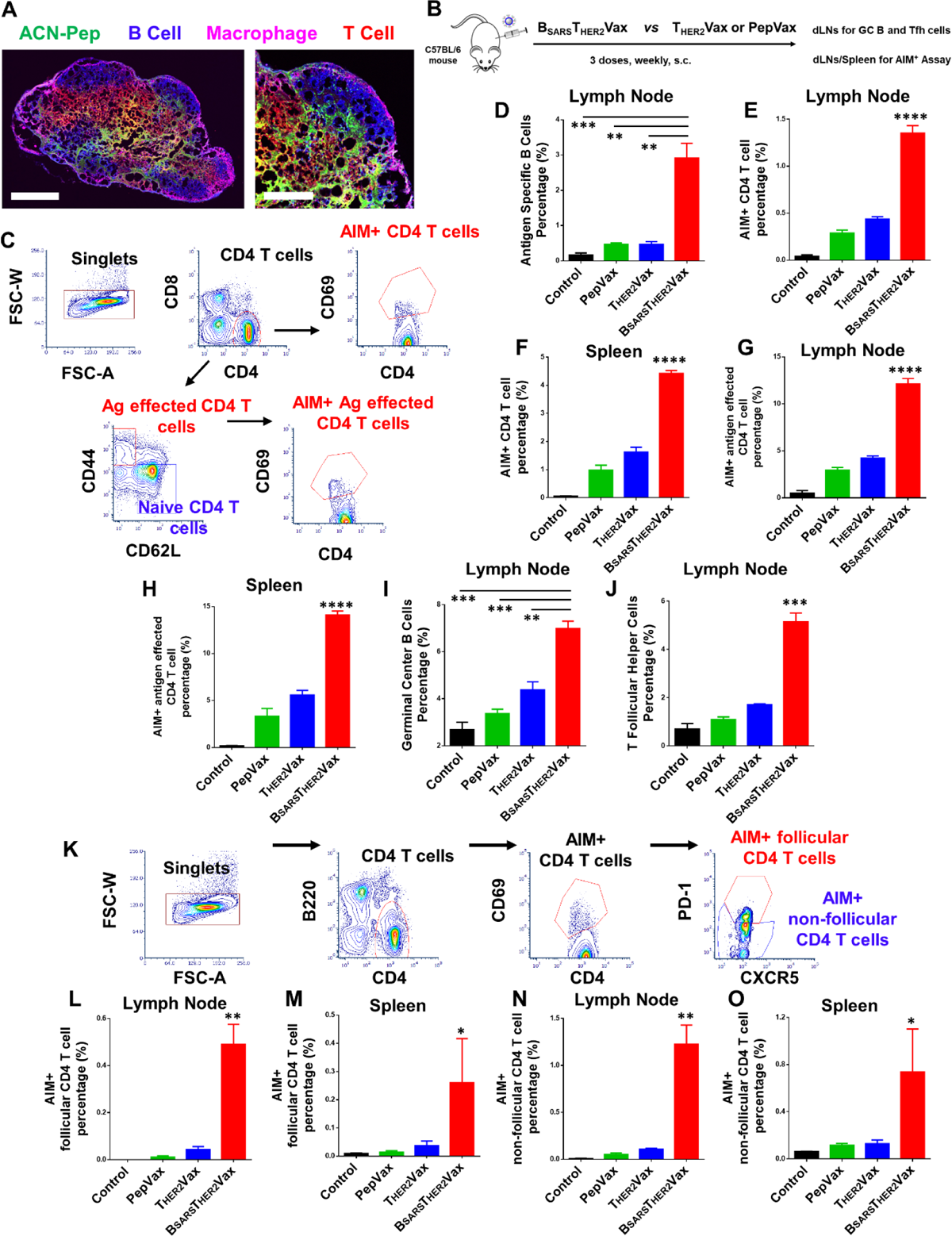
SARS-CoV-2 B cell epitope guided T cell NanoVaccines induced SARS-CoV-2 specific B cell and tumor specific CD4 T cell activation in vivo. (A) Confocal imaging of ACN penetration into lymph nodes. The scale bar is 200 µm in whole-lymph-node images and 100 µm in magnified images. (B) Schematic illustration of C57BL/6 mouse immunization with vaccines, ACN conjugated with SARS-CoV-2 B cell epitope and tumor HER2 CD4/CD8 T antigen (B_SARS_T_HER2_Vax), ACN conjugated with tumor HER2 CD4/CD8 T antigen alone (T_HER2_Vax), and tumor HER2 CD4/CD8 T antigen (PepVax). 50 μg peptide for each antigen with 10 μg 2’3’-cGAMP was used for three times weekly by subcutaneous injection. (C) Cells isolated from lymph nodes or spleens after vaccination were restimulated with HER2 CD4/CD8 epitopes, and measured activation-induced markers positive (AIM^+^) markers to identify antigen specific CD4 T cells. Gating strategy of AIM^+^ CD4 T cells (CD8^-^CD4^+^CD69^+^) and AIM^+^ antigen effected CD4 T cells (CD8^-^CD4^+^CD44^+^CD62L^-^CD69^+^). (D) Flow cytometry quantification of SARS-COV-2 antigen specific B cells (CD3^-^B220^+^Tetramer^+^) in lymph node. (E-G) Flow cytometry quantification of AIM^+^ CD4 T cells in lymph node (E) and spleen (F) and AIM^+^ antigen effected CD4 T cells in lymph node (G) and spleen (H). (I, J) Flow cytometry quantification of germinal center (GC, CD3^-^ B220^+^CD95^+^GL-7^+^) B cells (I) and T follicular helper (Tfh, B220^-^CD4^+^CXCR5^+^PD-1^+^) cells (J) from lymph node. (K) Gating strategy of AIM^+^ follicular CD4 T cells (AIM^+^CXCR5^+^PD-1^+^) and AIM^+^ non-follicular CD4 T cells (AIM^+^CXCR5^-^PD-1^-^). (L-O) Flow cytometry quantification of AIM^+^ follicular CD4 T cells in lymph node (L) and spleen (M), and AIM^+^ non-follicular CD4 T cells in lymph node (N) and spleen (O). Data for quantification are shown as mean ± SD, n = 3. For **Figure 2D-2O**, statistical comparisons were conducted between B_SARS_T_OT-II_ group with other groups. Statistical comparisons are based on one-way ANOVA, followed by post hoc Tukey’s pairwise comparisons or by Student’s unpaired T-test. The asterisks denote statistical significance at the level of * p < 0.05, ** p < 0.01, *** p < 0.001, **** p < 0.0001. ANOVA, analysis of variance; SD, standard deviation; n.s., no statistical significance.

To study whether B_SARS_T_tumor_Vax could induce B cell antigen presentation, resulting a higher level of SARS-CoV-2 specific B cells and tumor specific CD4 T cells in vivo, we immunized the mice with vaccines for three times using ACN conjugated with SARS-CoV-2 B epitope and human tumor HER2 CD4/CD8 T cell epitope (B_SARS_T_HER2_Vax) or human HER2 CD4/CD8 T cell epitope alone (T_HER2_Vax). The immune cells were monitored 10 days after the third immunization (**Figure 2B**). The results indicated that B_SARS_T_HER2_Vax stimulated 6.2-fold higher SARS-specific B cells (2.9% of all B cells, **Figure 2D**). We further employed an activation-induced marker (AIM) assay to identify the activation of antigen specific CD4 T cells (**Figure 2C**).^70,71^ B_SARS_T_HER2_Vax stimulated 3.1-fold and 2.7-fold higher activation of total antigen specific CD4 T cells than T_HER2_Vax in lymph node (1.35% vs 0.44%, **Figure 2E**) and spleen (4.42% vs 1.63%, **Figure 2F**), respectively. Among antigen specific CD4 T cells, B_SARS_T_HER2_Vax induced robust antigen specific antigen effected CD4 T cells compared to T_HER2_Vax in lymph node (12.10% vs 4.22%, **Figure 2G**) and spleen (14.11% vs 5.60%, **Figure 2H**).

In classical B cell activation, B cells present antigens to CD4 T cells, further activating the T follicular helper (Tfh) cell-dependent germinal center (GC) response. GC B cell and Tfh cell activation in tumors, especially in tertiary lymphoid structures (TLSs), is correlated with better outcomes after anti-PD-1/PD-L1 immunotherapy.^38,45,47,49,72–75^ Therefore, to study whether B_SARS_T_HER2_Vax induces GC-Tfh response which typically follows the B and CD4 T cell crosstalk, we monitored the germinal center (GC) B cell, Tfh cells after the above treatment. (**Figure 2B**). The results indicated that B_SARS_T_HER2_Vax stimulated 1.6-fold higher GC B cells (7.0% of all B cells, **Figure 2I**) and 3.5-fold higher T follicular helper cells (5.15% out of all CD4 T cells, **Figure 2J**) compared to T_HER2_Vax (0.47% antigen specific B cells, 4.4% GC B cells, 0.9% antigen specific GC B cells and 1.72% of Tfh cells). Moreover, B_SARS_T_HER2_Vax induced an 11-fold (0.49% vs 0.04% among total CD4 T cells) and 7-fold higher (0.26% vs 0.04% among total CD4 T cells) higher antigen specific Tfh cells compared to T_HER2_Vax in lymph node and spleen, respectively (**Figure 2K-2M**).

Interestingly, we found that B_SARS_T_HER2_Vax also induced a 11.4-fold (1.23% vs 0.11% among total CD4 T cells) and 5.8-fold (0.74% vs 0.13% among total CD4 T cells) higher antigen specific non-follicular CD4 T cell compared to T_HER2_Vax group in lymph node and spleen, respectively (**Figure 2N** and **2O**). These results indicated that SARS-CoV-2 B epitope-guided cancer vaccines activated two separate CD4 T cell subpopulations followed by SARS-CoV-2 specific B cells present antigen to tumor-specific CD4 T cells. In accordance with classic B cell immunity, B and CD 4 T cell crosstalk activates SARS-specific geminal center (GC) B cells and tumor-specific follicular T helper cells (Tfh) in the germinal center. However, we also observed that B/CD4 T cell crosstalk activates tumor-specific non-follicular CD4 T cell immunity (non-Tfh) outside germinal center.

### SARS-CoV-2 B epitope-guided neoantigen NanoVaccines enhanced anticancer efficacy compared to neoantigen vaccine without B cell antigen presentation function

To study if SARS-CoV-2 B epitope-guided cancer vaccines (B_SARS_T_tumor_Vax) could enhance anticancer efficacy, we used four different cancer models (breast cancer, OVA-melanoma, melanoma, and pancreatic cancer). We generated four types of vaccines by conjugating ACN with SARS-CoV-2 B epitopes and tumor antigens, which include HER2^+^ breast cancer (tumor HER2 antigen), B16-OVA (OVA antigen, MHC-I-restricted OVA_257-264_ peptide and MHC-II-restricted OVA_323-339_), melanoma (neoantigen, MHC-I-restricted M27^63,64^, and MHC-II-restricted M30), pancreatic cancer (neoantigen, Kras G12D, and Tp53 R172H)^63,64^. These four types of vaccines were administered to the above four different tumor mouse models by subcutaneous injection (SC) for three doses. In HER2^+^ breast cancer mouse model using D2E2/F2 cells, B_SARS_T_HER2_Vax, combined with an anti-PD-1 antibody, significantly inhibited tumor growth by 88.7 %. In contrast, the ACN vaccine conjugated with only the human HER2 CD4/CD8 T cell epitope (T_HER2_Vax) showed less anticancer efficacy (**Figure 3A and S5**). In melanoma B16-OVA model, SARS-CoV-2 B epitope-guided cancer vaccine (B_SARS_T_OVA_Vax), combined with an anti-PD-1 antibody, significantly inhibited tumor growth by 96.5%, which is more efficacious than T_OVA_Vax where ACN was conjugated with OVA T antigen alone (**Figure 3B**).

**Fig. 3.**
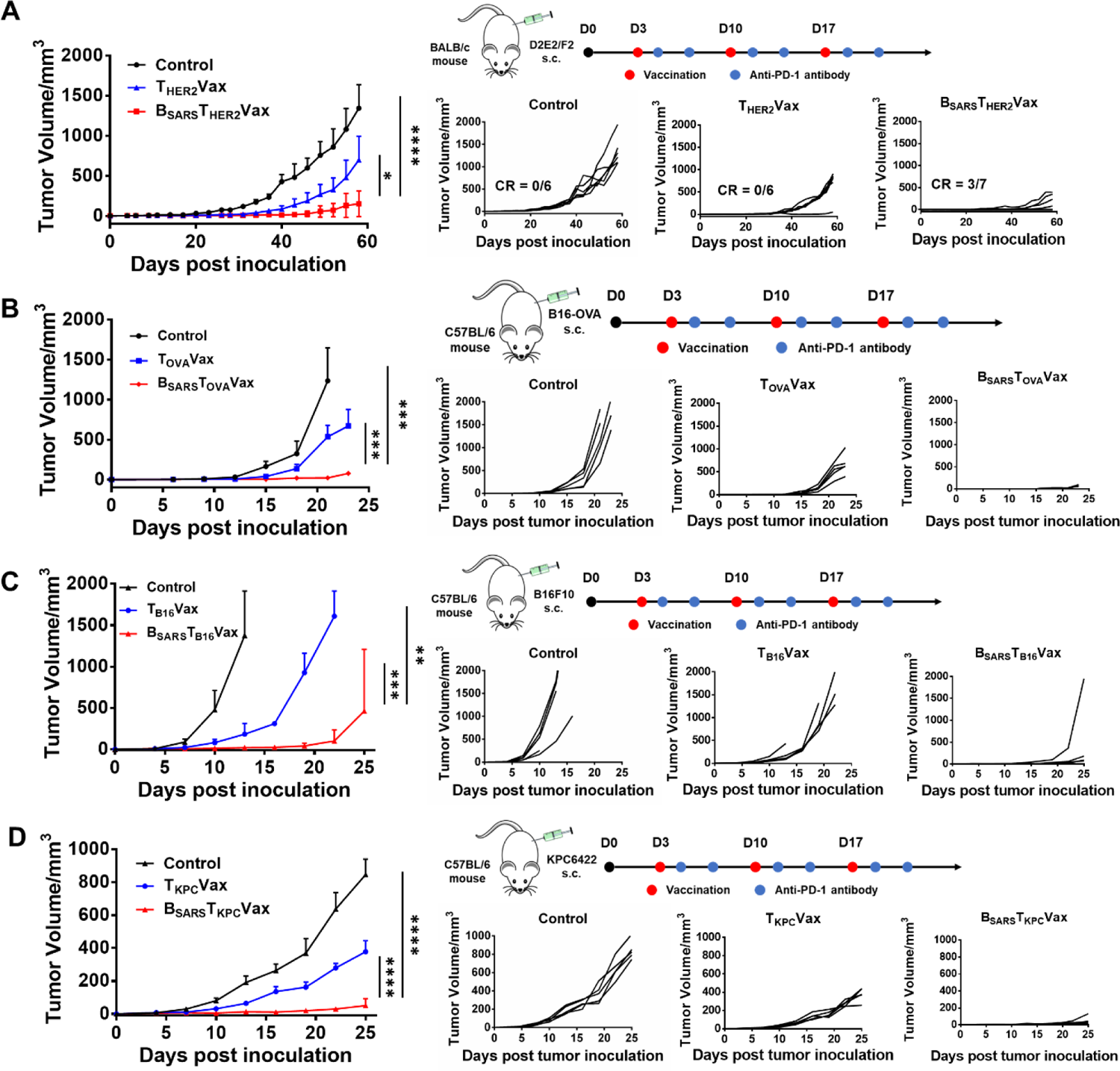
SARS-CoV-2 B-epitope guided neoantigen cancer NanoVaccines enhanced anticancer efficacy compared to neoantigen vaccine without B cell antigen presentation function(A) Antitumor efficacy of SARS-CoV-2 B epitope and tumor HER2 CD4/CD8 T antigen (B_SARS_T_HER2_Vax) compared with the vaccine with tumor HER2 CD4/CD8 T antigen (T_HER2_ Vax) in breast cancer model with subcutaneously inoculated with D2E2F2 cells. (B) Anticancer efficacy of SARS-CoV-2 B epitope-guided cancer vaccine B_SARS_T_OVA_Vax in a melanoma B16-OVA model compared to vaccine with T cell antigen alone (T_OVA_Vax). (C) Antitumor efficacy of SARS-CoV-2 B epitope-guided neoantigen cancer vaccine (B_SARS_T_B16_Vax) in B16 melanoma model in comparison with vaccine with T neoantigen alone (T_B16_ Vax). (D) Antitumor efficacy of SARS-CoV-2 B epitope neoantigen cancer vaccine (B_SARS_T_KPC_ Vax) in pancreatic cancer model in comparison with vaccine with T cell neoantigen alone (T_KPC_ Vax). The mice were subcutaneously inoculated with D2E2F2 breast cancer (2.5 × 10^5^) cells, or B16-OVA (2 × 10^5^) cells, or melanoma B16F10 (5 × 10^5^) cells, or pancreatic cancer KPC6422 (5 × 10^5^) cells, respectively. Then mice were vaccinated with various vaccines (50 μg peptide for each antigen with 10 μg 2’3’-cGAMP, subcutaneous injection) for three times weekly. Anti-PD-1 antibody (100 μg) was intraperitoneally administrated to mice. Tumor growth was monitored every three days. Data for quantification are shown as mean ± SD, n = 5. Statistical comparisons are based on one-way ANOVA, followed by post hoc Tukey’s pairwise comparisons or by Student’s unpaired T-test. The asterisks denote statistical significance at the level of * p < 0.05, ** p < 0.01, *** p < 0.001, **** p < 0.0001. ANOVA, analysis of variance; SD, standard deviation; n.s., no statistical significance.

In B16F10 melanoma model, SARS-CoV-2 B epitope-guided neoantigen cancer vaccine (B_SARS_T_B16_Vax) enhanced the antitumor efficacy (98.6% tumor growth inhibition) than T_B16_Vax where ACN was conjugated with only T cell neoantigen alone (T_B16_Vax) (**Figure 3C**). In pancreatic cancer model using KPC6422 pancreatic cancer cells, SARS-CoV-2 B epitope-guided neoantigen cancer vaccine (B_SARS_T_KPC_Vax) substantially inhibited tumor by 94.0% (**Figure 3D**) compared to the vaccine where ACN was conjugated with T cell neoantigen alone (T_KPC_Vax). These data clearly showed that SARS-CoV-2 B epitope-guided cancer vaccines (B_SARS_T_tumor_Vax) could enhance anticancer efficacy in four different types of cancer mouse models compared to vaccines without B cell antigen presentation function.

### SARS-CoV-2 B epitope-guided neoantigen NanoVaccines enhanced GC/Tfh responses and activated tumor specific CD4 T cells in four different cancer mouse models

To further assess if these vaccines enhanced the GC/Tfh responses and activated tumor-specific CD 4 T cell in all four types of cancer as described above, we performed the flow cytometry analysis of the B and CD 4 T cell phenotypes (**Figure 4A**). In HER2+ breast cancer, B_SARS_T_HER2_ Vax increased 2.4-fold GC B cells (14.2% in lymph node among B cells) and 3.9-fold Tfh cells (8.4% in lymph node among CD4 T cells) compared to T_HER2_Vax (6.0% GC in lymph node, 2.2% Tfh in lymph node) (**Figure 4B**). In melanoma B16-OVA mouse model, B_SARS_T_OVA_Vax increased 2-fold GC B cells (18.29% in lymph node among B cells), 6.9-fold Tfh cells (6.31% in lymph node among CD4 T cells) and 4-fold tumor antigen specific CD4 T cells (15.73% in lymph node among CD4 T cells) compared to T_HER2_ Vax (9.22% GC B cells, 0.92% Tfh cells and 3.88% antigen specific CD4 T cells) (**Figure 4C**).

**Fig. 4.**
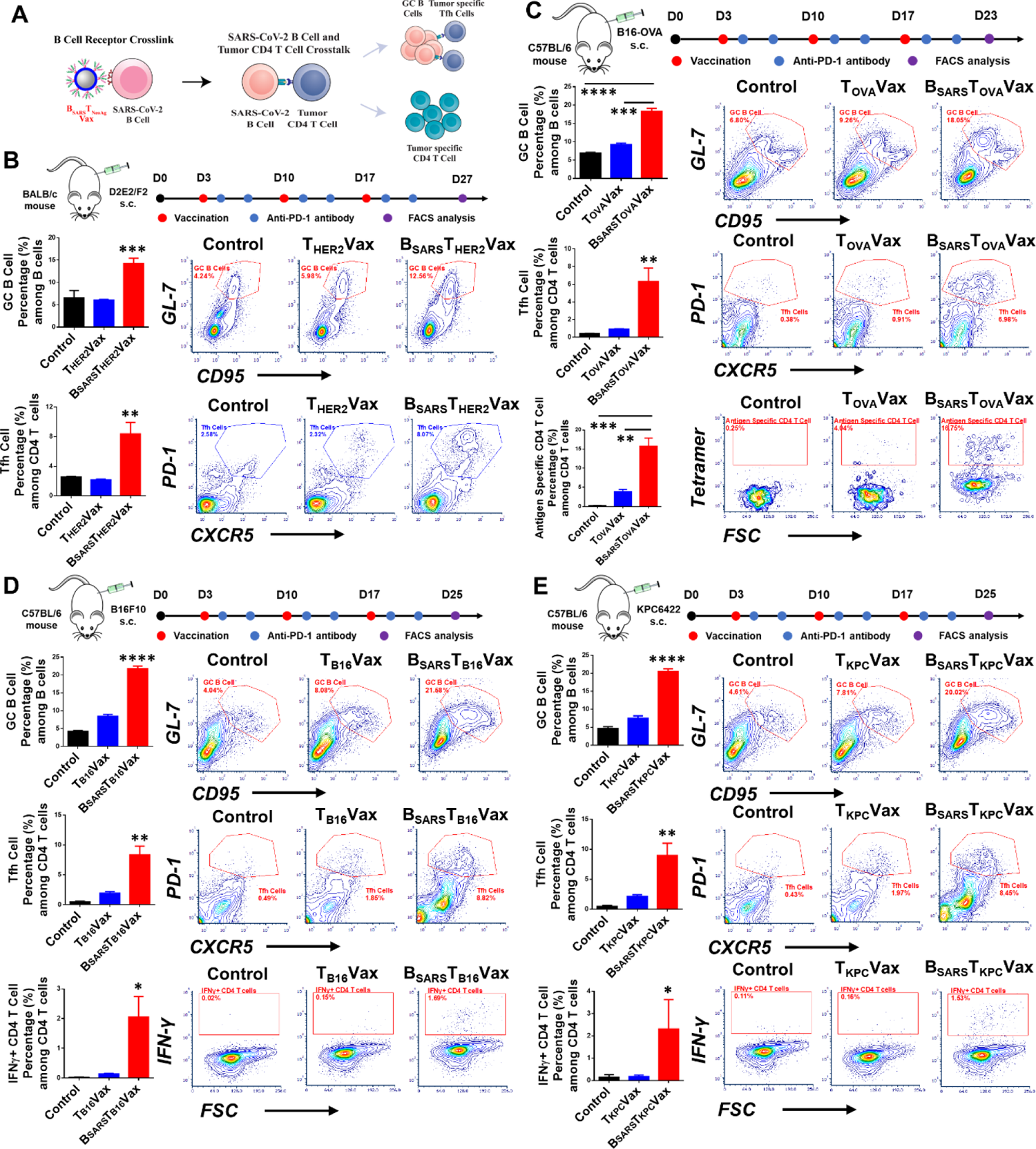
SARS-CoV-2 B epitope-guided neoantigen NanoVaccines enhanced GC/Tfh responses and elicited tumor-specific CD4 T cell. (**A**) Schematic illustration of activation of germinal center (GC) B/Tfh responses and tumor antigen specific CD4 T cells by SARS-CoV-2 B epitope-guided neoantigen vaccine (B_SARS_T_NeoAg_Vax). (**B**) Flow cytometry quantification and representative analysis of GC B cells (B220^+^CD95^+^GL-7^+^) and Tfh cells (B220^-^CD4^+^CXCR5^+^ PD-1^+^) from lymph node 10 days after three vaccinations in D2E2/F2 HER2+ breast cancer model. Mice samples are obtained from **Figure 3A**. (**C**) Flow cytometry quantification and representative analysis of lymph node GC B cells, Tfh cells and tumor OT-II specific CD4 T cells (CD45^+^CD8^-^CD4^+^Tetramer^+^) from melanoma B16-OVA model (mice samples are obtained from **Figure 3B**, 6 days after the third vaccination). (**D**) Flow cytometry quantification and representative analysis of lymph node GC B cells, Tfh cells and spleen B16-F10 neoantigen specific CD4 T cells (CD45^+^CD8^-^CD4^+^IFN-γ^+^) from melanoma B16-F10 model (mice samples are obtained from **Figure 3C**, 8 days after the third vaccination). (**E**) Flow cytometry quantification and representative analysis of lymph node GC B cells, Tfh cells and spleen KPC neoantigen specific CD4 T cells (CD45^+^CD8^-^CD4^+^IFN-γ^+^) from pancreatic KPC model (mice samples are obtained from **Figure 3D**, 8 days after the third vaccination). Data for quantification are shown as mean ± SD, n = 3. Statistical comparisons were conducted between B_SARS_T_NeoAg_ group with other groups. Statistical comparisons are based on one-way ANOVA, followed by post hoc Tukey’s pairwise comparisons or by Student’s unpaired T-test. The asterisks denote statistical significance at the level of * p < 0.05, ** p < 0.01, *** p < 0.001, **** p < 0.0001. ANOVA, analysis of variance; SD, standard deviation; n.s., no statistical significance.

In B16F10 melanoma tumor model, B_SARS_T_B16_Vax elicited 2.6-fold of GC B cells, 4.3-fold of Tfh cells and 14.7-fold of tumor antigen specific CD4 T cells than T_B16_Vax (**Figure 4D**). In pancreatic cancer model using KPC6422, B_SARS_T_KPC_Vax activated 2.7-fold of GC B cells and 4.2-fold of Tfh cells than T_KPC_Vax in tumor (**Figure 4E**). B_SARS_T_KPC_Vax also elicited 13-fold of tumor antigen specific CD4 T cells (2.31%) compared to T_KPC_Vax (0.18%) in tumor (**Figure 4E**). In the KPC6422 pancreatic cancer model, we further used single cell analysis to monitor CD 4 T cell phenotypes in tumors post-vaccination. B_SARS_T_KPC_Vax increased cytotoxic CD4 T cell (Gzmb) and Tfh cell (Cxcr5), while decreased exhausted CD4 T cell (Pdcd1) and CD4 T regulatory cells (Foxp3) compared to T_KPC_Vax (**Figure S6**). In summary, SARS-CoV-2 B epitope-guided cancer vaccine enhanced GC and Tfh interaction, and elicited tumor specific CD4 T cell in all four cancer models.

### The efficacy of SARS-CoV-2 B epitope-guided cancer NanoVaccine is dependent on B and CD4 T cell activation

To confirm that the improved antitumor efficacy by SARS-CoV-2 B epitope-guided cancer NanoVaccine is B cell dependent, we evaluated its efficacy on µMT mice (**Figure S7**, lacking mature B cells) bearing B16-OVA melanoma. The results showed that B_SARS_T_OVA_Vax failed to inhibit tumor growth in µMT mice compared to the control group without treatment (**Figure 5A and S8A**). However, infusion of B cells isolated from the spleen of wild type mice (C57BL/6) into µMT mice restored the effectiveness of B_SARS_T_OVA_Vax, which is comparable to wild type mice immunized with B_SARS_T_OVA_Vax (**Figure 5A and S8A**). These data suggest that the efficacy of B_SARS_T_OVA_Vax is B cell dependent. Furthermore, flow cytometry analysis revealed that B_SARS_T_OVA_ Vax did not induce GC B cells or Tfh cells, as well as increased much less antigen specific CD4 T cells in µMT mice. Infusion of B cells into µMT mice significantly boosted GC B cells, Tfh cells, and antigen specific CD4 T cells after B_SARS_T_OVA_Vax vaccination (**Figure 5B, and S8B**). These findings suggest that the activation of tumor antigen-specific CD4 T cell by B_SARS_T_OVA_Vax is B cell dependent.

**Fig. 5.**
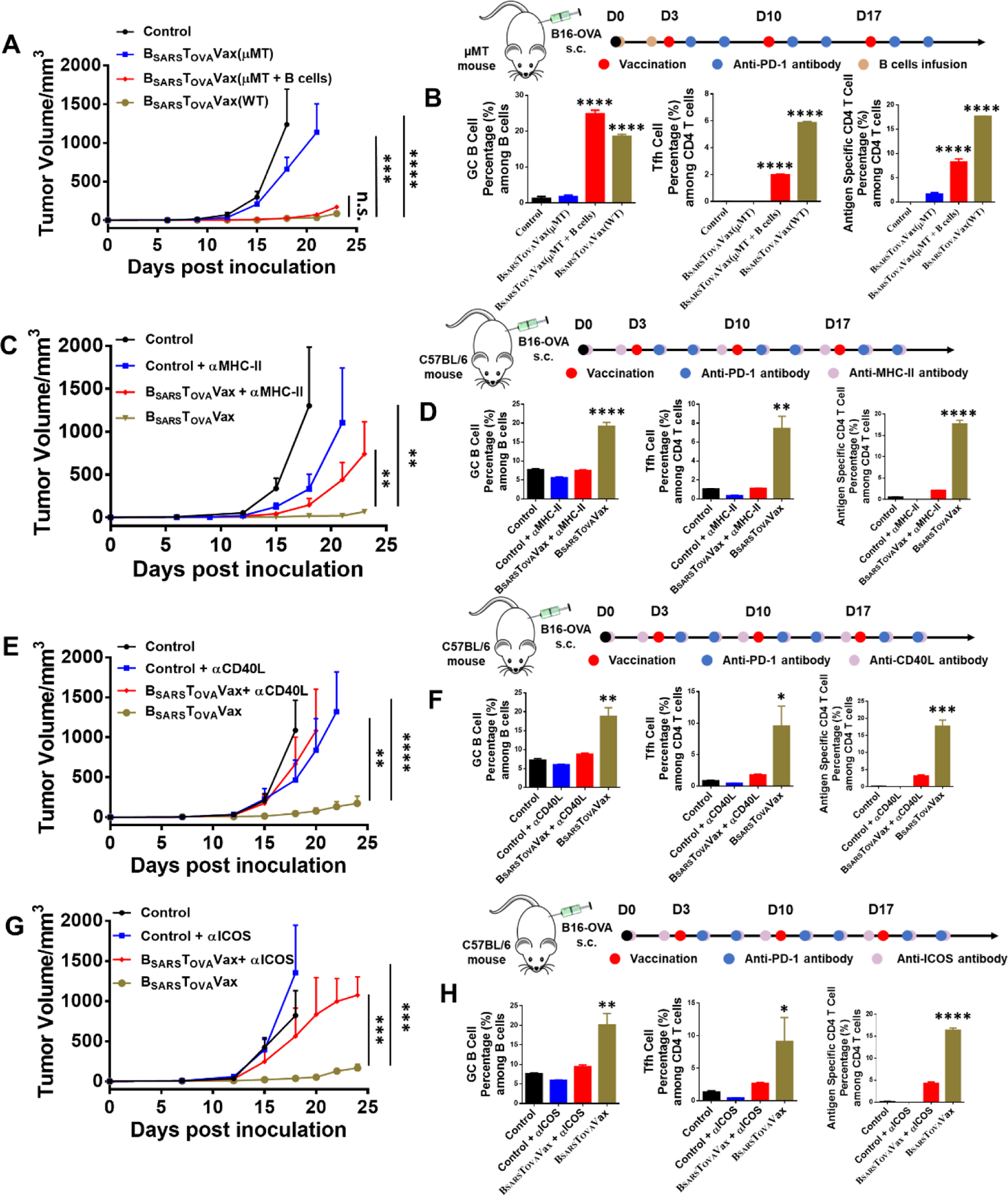
The anticancer efficacy of SARS-CoV-2 B cell epitope-guided cancer NanoVaccine is dependent on B cell and CD4 T cell activation. (A) Antitumor efficacy of SARS-CoV-2 B epitope-guided cancer vaccine (B_SARS_T_OVA_Vax) in µMT mice (without mature B cells) bearing B16-OVA melanoma after three vaccinations in comparison to vaccine with T cell antigen alone. In B cell infusion group, 2 × 10^6^ B cells isolated from male C57BL/6 mice were infused to male µMt mice 1 day and 3 days before immunization to restore their B cell function for anticancer efficacy. (B) Flow cytometry quantification of GC B cells (B220^+^CD95^+^GL-7^+^) and Tfh cells (B220^-^CD4^+^CXCR5^+^PD-1^+^) from lymph node and OT-II specific CD4 T cells (CD45^+^CD8^-^ CD4^+^Tetramer^+^) from tumor 6 days after the third vaccinations from (A). (C) Antitumor efficacy of B_SARS_T_OVA_Vax in B16-OVA melanoma mice treated with or without αMHC-II antibody. αMHC-II antibody (500 µg/mouse) was intraperitoneal injected 3 days before vaccination, and then dosed 200 µg every two days. (D) Flow cytometry quantification of GC B cells and Tfh cells from lymph node and OT-II specific CD4 T cells from tumor 6 days after the third vaccination from (C). (E) Antitumor efficacy of B_SARS_T_OVA_Vax in B16-OVA melanoma mice treated with or without αCD40L antibody. αCD40L antibody (200 µg/mouse) was intraperitoneal injected 3 days before vaccination, and then dosed 200 µg every two days. (F) Flow cytometry quantification of GC B cells and Tfh cells from lymph node and OT-II specific CD4 T cells from tumor 6 days after the third vaccinations from (E). (G) Antitumor efficacy of B_SARS_T_OVA_Vax in B16-OVA melanoma mice with or without αICOS antibody. αICOS antibody (200 µg/mouse) was intraperitoneal injected 3 days before vaccination, and then dosed 200 µg every two days. (H) Flow cytometry quantification of GC B cells and Tfh cells from lymph node and OT-II specific CD4 T cells from tumor 6 days after the third vaccinations from (G). For vaccinations in **Figure 5A**, 5C, 5E and 5G, 50 μg peptide for each antigen with 10 μg 2’3’-cGAMP, was used by subcutaneous injection, 100 μg of anti-PD-1 was intraperitoneally administrated to mice. For **Figure 5A**, B_SARS_T_OVA_Vax group is compared with all other groups. For **Figure 5B**, B_SARS_T_OVA_Vax group and B_SARS_T_OVA_Vax (µMT + B cells) group are compared with all other groups. For **Figure 5C-5H**, B_SARS_T_OVA_Vax group is compared with control and B_SARS_T_OVA_Vax plus αMHC-II, αCD40L or αICOS groups. Statistical comparisons are based on one-way ANOVA, followed by post hoc Tukey’s pairwise comparisons or by Student’s unpaired T-test. The asterisks denote statistical significance at the level of * p < 0.05, ** p < 0.01, *** p < 0.001, **** p < 0.0001. ANOVA, analysis of variance; SD, standard deviation; n.s., no statistical significance.

To verify that the enhanced antitumor efficacy of the SARS-CoV-2 B epitope-guided cancer NanoVaccines also relies on CD 4 T cell activation, we assessed the vaccine’s antitumor efficacy in mice with B16-OVA melanoma, using αMHC-II, αCD40L or αICOS to inhibit CD4 T cell activation. The anticancer efficacy of B_SARS_T_OVA_Vax was significantly reduced when mice were treated with αMHC-II (**Figure 5C and S9A**), αCD40L (**Figure 5E and S10A**) or αICOS (**Figure 5G and S11A**) antibodies. In addition, these antibodies also considerably inhibited the activation of GC B cells, Tfh cells, and antigen specific CD4 T cells (**Figure 5D, 5F, 5H, S9B, S10B and S11B**).

To further confirm the enhanced antitumor efficacy of the SARS-CoV-2 B epitope-guided cancer NanoVaccine is CD 4 T cell activation dependent, we also evaluated the vaccine’s antitumor efficacy in D2E2/F2 HER2+ breast cancer model using αCD40L antibody to inhibit CD4 T cell activation. αCD40L antibody significantly reduced the anti-cancer efficacy of B_SARS_T_HER2_Vax vaccination (**Figure S12**). Flow cytometry analysis showed that αCD40L blockage reduced GC B cells and Tfh cells in lymph nodes (**Figure S13**). In summary, these results indicated that B cell and CD4 T cell activation is essential for the effectiveness of the SARS-CoV-2 B epitope-guided cancer vaccine.

### SARS-CoV-2 B Epitope-Guided Cancer NanoVaccines Induce Tumor Antigen-Specific CD8 T Cell Response Through B Cell and CD4 T Cell Activation

To evaluate how the SARS-CoV-2 B epitope-guided cancer NanoVaccine affects CD 8 T cell immunity, we used single cell analysis to monitor CD 8 T cell phenotypes in tumors post-vaccination. Interestingly, B_SARS_T_KPC_Vax also induced a higher percentage of cytotoxic CD8 T cells (highly expressed Prf1), while reducing the percentage of CD8 T regulatory cells (highly expressed Foxp3) and exhausted CD8 T cell (highly expressed Pdcd1, Eomes and Lag3 genes) than T_KPC_ Vax (**Figure 6A, S14A**). In contrast, the neoantigen vaccine T_KPC_ Vax with CD4/CD8 T cells antigen did not reduce CD8 T regulatory cell and exhausted CD8 T cells, which was in agreement with previous clinical studies using neoantigen vaccines in melanoma patients^76,77^.

**Fig. 6.**
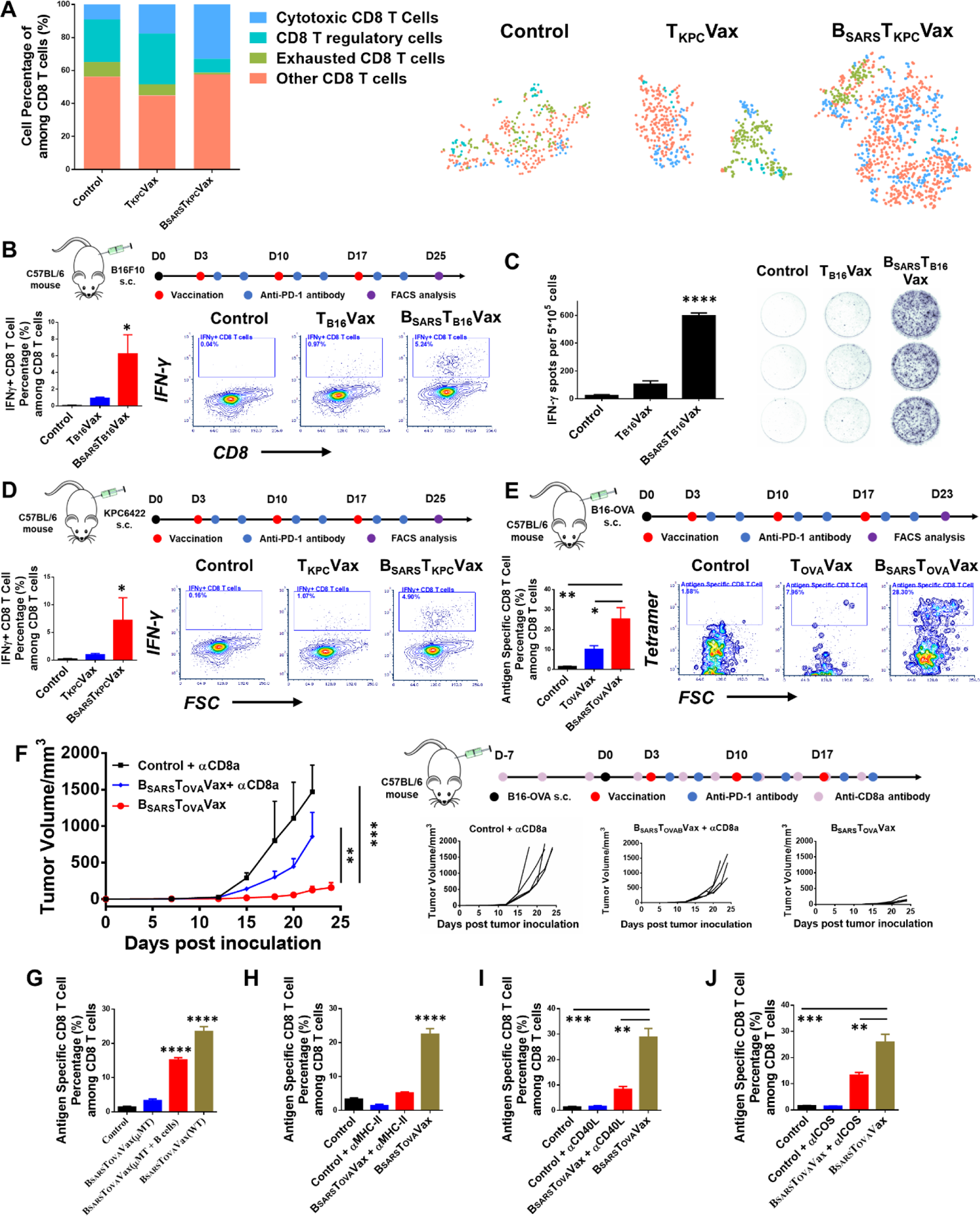
SARS-CoV-2 B Epitope-Guided Cancer NanoVaccines Induce Tumor Antigen-Specific CD8 T Cell Response Through B Cell and CD4 T Cell Activation. (A) Stacked bar charts and t-SNE plot of CD8 T cells from tumor-infiltrating immune cells. (B) Flow cytometry quantification and representative analysis of spleen B16-F10 neoantigen specific CD8 T cells (CD45^+^CD4^-^CD8^+^IFN-γ^+^) from B16-F10 melanoma model (mice samples are obtained from **Figure 3C**, 8 days after the third vaccination). (C) ELISPOT analysis of IFN-spot-forming cells among splenocytes after *ex vivo* restimulation with B16-F10 neoantigens on day 25 from **Figure 3C**. (D) Flow cytometry quantification and representative analysis of spleen KPC neoantigen specific CD8 T cells (CD45^+^CD4^-^CD8^+^IFN-γ^+^) from KPC pancreatic cancer model (mice samples are obtained from **Figure 3D**, 8 days after the third vaccination). (E) Flow cytometry quantification and representative analysis of tumor OT-I specific CD8 T cells (CD45^+^CD4^-^ CD8^+^Tetramer^+^) from B16-OVA melanoma model (mice samples are obtained from **Figure 3B**, 6 days after the third vaccination). Data for quantification are shown as mean ± SD, n = 3. (F) Antitumor efficacy of SARS-CoV-2 B epitope-guided cancer vaccine (B_SARS_T_OVA_Vax) in B16-OVA melanoma mice treated with or without αCD8a antibody. αCD8a antibody (200 µg/mouse) was intraperitoneal injected 10 days before vaccination, and then dosed 200 µg every three days. (G-J) Flow cytometry quantification of tumor OT-I specific CD8 T cells from tumor 6 days after the third vaccination (G from **Figure 5A**, H from **Figure 5C**, I from **Figure 5E**, J from **Figure 5G**). For **Figure 6B-6F**, B_SARS_T_OVA_ group is compared with all other groups. For **Figure 6G**, B_SARS_T_OVA_ group and B_SARS_T_OVA_ (µMT with B cells infusion) group are compared with all other groups. For **Figure 6H-6J**, B_SARS_T_OVA_ group is compared with the control and groups of B_SARS_T_OVA_ plus αMHC-II, αCD40L or αICOS antibody. Statistical comparisons are based on one-way ANOVA, followed by post hoc Tukey’s pairwise comparisons or by Student’s unpaired T-test. The asterisks denote statistical significance at the level of * p < 0.05, ** p < 0.01, *** p < 0.001, **** p < 0.0001. ANOVA, analysis of variance; SD, standard deviation; n.s., no statistical significance.

To confirm if SARS-CoV-2 B epitope-guided cancer NanoVaccine promoted tumor antigen specific CD8 T cell activation, we used flow cytometry and ELISPOT to measure the antigen specific CD8 T cells after vaccination in the B16F10 melanoma, KPC6422 pancreatic cancer and B16-OVA melanoma mouse models. In B16F10 melanoma model, B_SARS_T_B16_Vax dramatically induced higher antigen specific cytotoxic CD8 T cells by flow cytometry (6.8-fold) and by ELISPOT (5.7-fold) compared to T_B16_Vax (**Figure 6B** and **6C**). In KPC6422 pancreatic cancer model, B_SARS_T_KPC_Vax immunization elicited 7.5-fold of antigen specific cytotoxic CD8 T cells compared to T_KPC_Vax (**Figure 6D and S14B**). In B16-OVA model, B_SARS_T_OVA_Vax stimulated 2.5-folder higher antigen specific cytotoxic CD8 T cell compared to T_OVA_Vax (**Figure 6E**).

To confirm whether the anticancer efficacy of SARS-CoV-2 B epitope-guided cancer NanoVaccine is dependent on CD 8 T cell, we assessed its efficacy of B_SARS_T_OVA_Vax in the B16-OVA melanoma model following CD8 T cell depletion using an αCD8 antibody. The results showed that the anticancer efficacy of B_SARS_T_OVA_Vax and tumor infiltrated antigen specific CD8 T cells was significantly reduced by the depletion of CD8 T cells with the αCD8 antibody (**Figure 6F**).

To determine whether the CD8 T cell immunity induced by the SARS-CoV-2 B epitope-guided cancer NanoVaccine is dependent on B cells, we analyzed the infiltration of antigen-specific CD8 T cells in B16-OVA melanoma tumors in both wild type and µMT mice after B_SARS_T_OVA_Vax vaccination. In µMT mice without mature B cells, the infiltration of antigen-specific CD8 T cells after vaccination was significantly reduced to 3.28% compared to 23.45% in wide type mice. However, infusion of normal B cells into µMT mice restored the infiltration of antigen-specific CD8 T cells in tumor to 15.19% (**Figure 6G and S15A**) after vaccination. These data suggested that CD8 activation by B_SARS_T_OVA_Vax is B cell dependent.

To further investigate whether the CD8 T cell immunity induced by the SARS-CoV-2 B epitope-guided cancer NanoVaccine is dependent on CD4 T cell activation, we assessed the antigen-specific CD8 T cells in tumors after vaccination in B16-OVA melanoma mice, using αMHC-II, αCD40L or αICOS to block CD4 T cell activation (**Figure 5**). The antigen-specific CD8 T cells in tumors after B_SARS_T_OVA_Vax vaccination were dramatically reduced when mice were treated with αMHC-II (5.15% vs. 22.48%, **Figure 6H and S15B**), αCD40L (8.25% vs. 28.82%, **Figure 6I and S15C**) or αICOS (13.14% vs. 25.88%, **Figure 6J and S15D**) antibodies. Altogether, these results showed that SARS-CoV-2 B epitope-guided cancer NanoVaccine promoted antigen specific CD8 T cell response, which is dependent on B cell and CD4 T cell activation.

### Develop SARS-CoV-2 B epitope-guided neoantigen mRNA cancer NanoVaccine for better anticancer efficacy in pancreatic cancer mouse model

To harness SARS-CoV-2 B epitope-guided strategies for mRNA neoantigen cancer NanoVaccines, we engineered a lipid nanoparticle (LNP) to encapsulate T cell mRNA neoantigens of P53 R172H and KRAS G12D (**Table S1**), where the LNP surface was conjugated with SARS-CoV-2 B peptide epitopes (B_SARS_T_KPC-mRNA_Vax) (**Figure 7A**). The mRNA sequences encoding peptides contain P53 R172H (MTEYKLVVVGADGVGKSALTIQLIQNH) and KRAS G12D mutations (IYKKSQHMTEVVRHCPHHERCSDGDGL). The lipid nanoparticle formulation includes 1,2-Distearoyl-sn-glycero-3-phosphocholine (DSPC), cholesterol, the specially designed ionizable lipid MMT6-55 (**Scheme 1**), and DMP-PEG-MAL for peptide conjugation (**Table S2**). We employed microfluidic techniques for encapsulating the mRNA neoantigens within the lipid nanoparticle, which was subsequently conjugated with the SARS-CoV-2 B peptide epitopes (SP14P5: CDDDTESNKKFLPFQQFGRDIA, SP21P2: CDDDPSKPSKRSFIEDLLFNKV), where the cysteines at its N-terminus was direct conjugated to the maleimide group on the surface of LNP. The optimized LNP formulation (N/P ratio of 14), with a size of 120 nm (**Figure 7B and Table S3**), demonstrated cellular transfection efficiency comparable to clinically used LNP formulations (**Figure 7C, S16A and S16B**). We encapsulated mRNA encoding KPC neoantigens (Kras G12D and Tp53 R172H) in the LNP, with or without surface-conjugated SARS-CoV-2 B cell epitope (B_SARS_T_KPC-mRNA_Vax and T_KPC-mRNA_Vax, respectively) (**Figure S16C**). The B_SARS_T_KPC-mRNA_Vax (20 μg mRNA, every four days for the first three vaccinations and then every three days for the last two vaccinations) significantly delayed tumor growth (**Figure 7D**). B_SARS_T_KPC-mRNA_Vax significantly enhanced GC B cells (**Figure 7E and S17A**), Tfh cells (**Figure 7F and S17A**), antigen-specific CD4 T cells (**Figure 7G and S17B**), and tumor antigen-specific CD8 T cell (**Figure 7H and S17B**) compared to the T_KPC-mRNA_Vax. Additionally, B_SARS_T_KPC-mRNA_Vax induced significantly higher levels of granzyme B+ CD4 T cells (**Figure 7I**) and granzyme B+ CD8 T cell (**Figure 7J**) than T_KPC-mRNA_Vax.

**Fig. 7.**
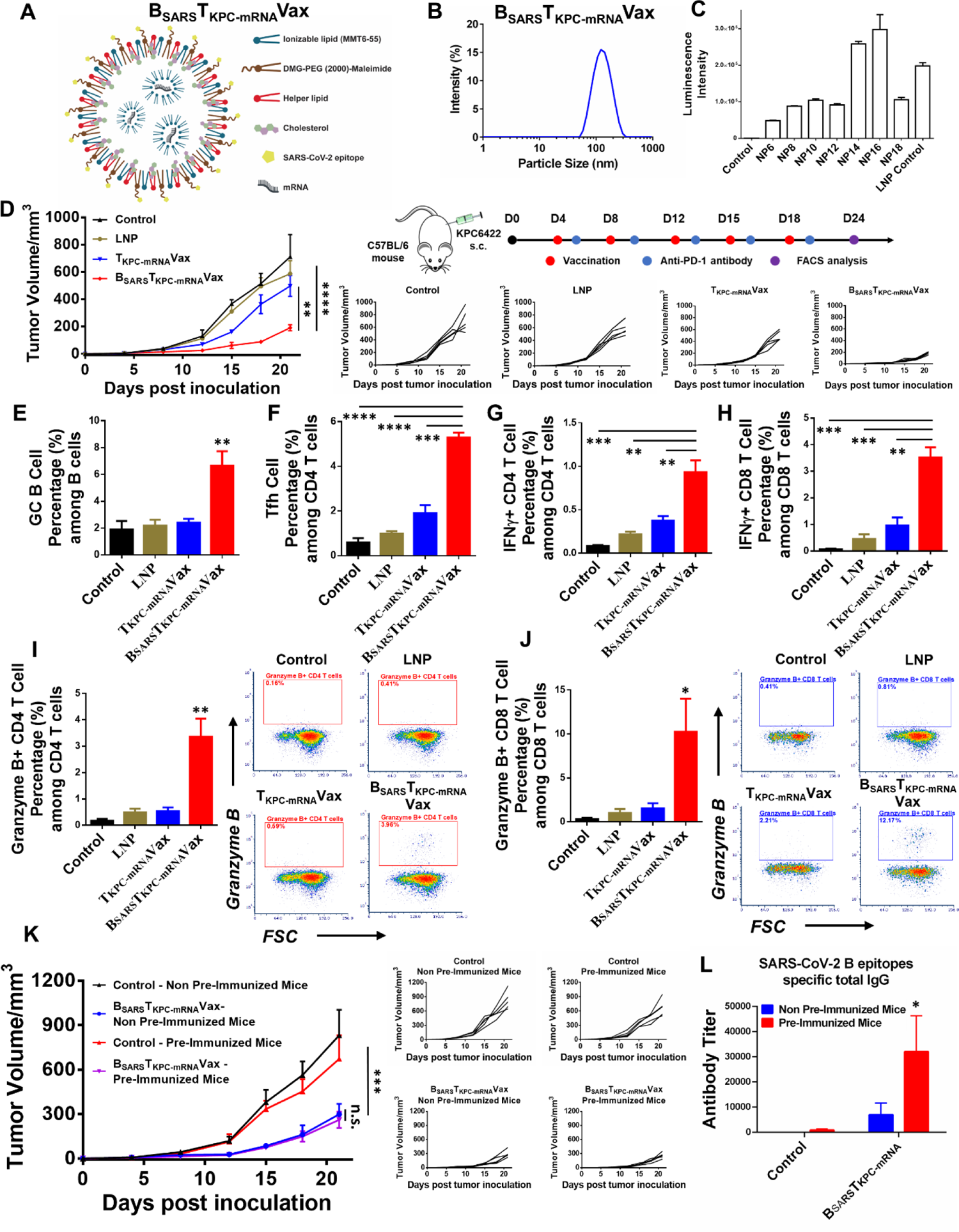
SARS-CoV-2 B epitope-guided neoantigen mRNA cancer NanoVaccine achieved better anticancer efficacy in pancreatic cancer model. (A) Schematic illustration SARS-CoV-2 B epitope-guided mRNA neoantigen cancer vaccines in a lipid nanoparticle (LNP) where its surface was conjugated with SARS-CoV-2 B epitope (B_SARS_T_KPC-mRNA_Vax). The mRNA sequences to encode peptide with Tp53 R172H and KRAS G12D mutations are listed in Table S2. (B) Size distribution of B_SARS_T_KPC-mRNA_Vax using DLS. (C) LNP formulation optimization based on the expression of firefly luciferase expression in DC2.4 cells transfected with firefly luciferase encoded mRNA encapsulated by SARS-CoV-2 B peptide conjugated LNP. The formulations were prepared with different ratio between the Amine groups (N) of the ionizable lipid and the Phosphate group (P) of mRNA (N/P ratio) and commercialized lipid nanoparticle (LNP Control) (D) Antitumor efficacy of B_SARS_T_KPC-mRNA_Vax compared with T_KPC-mRNA_Vax in pancreatic cancer model with KPC 6422 cells. (E-H) Flow cytometry quantification of lymph node GC B cells (B220^+^CD95^+^GL-7^+^) (E), lymph node Tfh cells (B220^-^CD4^+^CXCR5^+^PD-1^+^) (F), lymph node KPC neoantigen specific CD4 T cells (CD45^+^CD8^-^CD4^+^IFN-γ^+^) (G) and lymph node KPC neoantigen specific CD8 T cells (CD45^+^CD4^-^CD8^+^IFN-γ^+^) (H) 6 days after the final vaccination from (D). (I, J) Flow cytometry quantification and representative analysis of lymph node Granzyme B^+^ CD4 T cells (CD45^+^CD8^-^CD4^+^Granzyme^+^) (I) and lymph node Granzyme B^+^ CD8 T cells (CD45^+^CD4^-^CD8^+^Granzyme^+^) (J) 6 days after the final vaccination from (D). (K) Antitumor efficacy of B_SARS_T_KPC-mRNA_Vax in mice with or without pre-immunization using SARS-CoV-2 spike protein in KPC 6422 pancreatic cancer model. Mice are immunized with SARS-CoV-2 Spike ECD (5 µg, Val16-Pro1213, wild type, Alhydrogel® adjuvant 2%) every 7 days for 3 times to establish pre-existing SARS-CoV-2 immunity. (L) Quantification of antigen-specific IgG antibodies against SARS-CoV-2 B epitopes by indirect ELISA represented as antibody titer from serum collected 6 days after the final immunization. B_SARS_T_KPC-mRNA_VAX group is compared with all other groups. Statistical comparisons are based on one-way ANOVA, followed by post hoc Tukey’s pairwise comparisons or by Student’s unpaired T-test. The asterisks denote statistical significance at the level of * p < 0.05, ** p < 0.01, *** p < 0.001, **** p < 0.0001. ANOVA, analysis of variance; SD, standard deviation; n.s., no statistical significance.

Considering that many cancer patients may have pre-existing immunity against the SARS-CoV-2 spike protein, which could affect the effectiveness of SARS-CoV-2 B epitope-guided neoantigen cancer vaccines, we further compared the anticancer efficacy of B_SARS_T_KPC-mRNA_Vax in mice previously immunized with the SARS-CoV-2 spike protein (**Figure 7K**). The results indicated no significant difference in anticancer efficacy between mice with or without pre-existing immunity against the SARS-CoV-2 spike protein. Additionally, antibody responses against the SARS-CoV-2 epitopes were observed after three B_SARS_T_KPC-mRNA_Vax vaccinations in mice with or without previously immunized with the SARS-CoV-2 spike protein, where pre-immunized mice showed significantly higher antibody levels against SARS-CoV-2 B epitopes (**Figure 7L**).

## DISCUSSION

Cancer neoantigen vaccines typically rely on dendritic cell/macrophage-dependent antigen presentation to activate CD4/CD8 T cell antitumor immunity. These vaccines do not utilize B cell-mediated antigen presentation despite B cells being professional APCs.^24–26^ This exclusion is due to longstanding debates about the controversial role of B cell immunity in cancers.^27–35^ Emerging evidence suggested B cell-mediated antigen presentation also plays a vital role in anticancer immunotherapy.^36–44,78–80^ It is not known whether incorporating B cell-mediated antigen presentation into current neoantigen vaccines could enhance CD4 and CD8 T cell immunity to improve their anticancer efficacy.

In this study, we developed SARS-CoV-2 B cell epitope-guided neoantigen peptide or mRNA cancer vaccines (B_SARS_T_NeoAg_Vax or B_SARS_T_mRNA_Vax) to incorporate B cell antigen presentation function in neoantigen vaccine design. B_SARS_T_NeoAg_Vax promoted SARS-CoV-2 B cell-mediated antigen presentation, which not only activated classical GC response, but also enhanced tumor-specific follicular and non-follicular CD4 T cell immunity, as well as promoted B cell-dependent tumor-specific CD8 T cell immunity. Compared to current neoantigen cancer vaccines without the SARS-CoV-2 B epitope (T_NeoAg_Vax), B_SARS_T_NeoAg_Vax showed superior anticancer activity in melanoma, pancreatic, and breast cancer models. To implement this strategy in neoantigen mRNA cancer vaccine, we design a SARS-CoV-2 B epitope-guided neoantigen mRNA cancer vaccine (B_SARS_T_mRNA_Vax) using lipid nanoparticle (LNP), where its surface was modified with SARS-CoV-2 B peptide epitope, to encapsulate mRNA T cell neoantigens of P53 R172H and KRAS G12D. B_SARS_T_KPC-mRNA_Vax showed better anticancer efficacy, and enhanced tumor antigen specific CD4 and CD8 T cell immunity compared to conventional LNP mRNA neoantigen vaccine with T cell epitope that relies solely on DC/macrophage-mediated antigen presentation.

Our study presents a novel neoantigen vaccine design strategy that incorporates viral B cell mediated-antigen presentation to enhance tumor-specific CD4/CD8 T cell antitumor immunity. Compared to the current neoantigen cancer vaccine relying solely on DC/macrophage-mediated antigen presentation, B_SARS_T_NeoAg_Vax with peptide neoantigen enhanced 13- to 15-fold of tumor antigen specific CD4 T cells, and 6- to 8-fold of tumor antigen specific CD8 T cells in both B16F10 melanoma and KPC6422 pancreatic cancer models. B_SARS_T_mRNA_Vax with mRNA neoantigen enhanced 2.5-fold of tumor antigen specific CD4 T cells, and 3.7-fold of tumor antigen specific CD8 T cells. B_SARS_T_NeoAg_Vax activates SARS-CoV-2-specific B cells and expand these B cells for more effective antigen presentation.^24,25^ Our data aligns with previous studies showing that B cell-mediated antigen presentation benefits anticancer T cell immunity. For instance, TIL-B cells isolated from human tumors and tumor-draining lymph nodes can present tumor-associated antigens to directly activate CD4 T cells in vitro.^36,78^ Blocking B cell and CD4 T cell crosstalk (such as ICOS-ICOSL and CD40-CD40L interaction) impairs the anticancer immune responses and efficacy of immunotherapy in tumor-bearing mice.^79,80^ In addition, DCs and macrophages often become suppressed or dysfunctional in tumor immunosuppressive microenvironment, reducing their efficiency in antigen presentation.^81,82^ Therefore, utilizing B cells as antigen-presenting cells offers an alternative and effective strategy to enhance the efficacy of neoantigen vaccines.

It is worth noting that the underlying mechanisms of how B cell antigen presentation augments tumor specific CD4 and CD8 T cell immunity is not entirely clear and warrants further investigation. First, B cell antigen presentation-mediated B and CD4 T cell crosstalk is traditionally considered to be critical to activate GC response (GC B and Tfh cells) for antibody production.^72,74,75^ Indeed, there is direct evidence that tumor-infiltrated B cells secrete anti-tumor antibodies in situ.^72,83–86^ However, B_SARS_T_NeoAg_Vax enhanced anticancer efficacy through increased CD4 and CD8 T cell immunity, independent from antibody production in our study. Although B_SARS_T_NeoAg_Vax produced anti-SARS-CoV-2 antibodies, these antibodies do not contribute to its anticancer efficacy. These results align with previous findings that B cell depletion, but not plasma cell abrogation, enhances tumor growth in melanoma mouse model.^87^ Second, B_SARS_T_NeoAg_Vax not only promoted CG and Tfh response, but also activated a significant portion of tumor specific non-follicular CD4 T cells (>70%), which may directly combat the tumors.^88–90^ Previous studies showed that GC/Tfh response in tumors is correlated with better outcomes after anti-PD-1/PD-L1 immunotherapy.^38,45,47,49,72–75^ The function of non-follicular CD4 T cells through B cell-mediated antigen presentation warrants further investigation for their beneficial effects in cancer immunotherapy. Third, we also found B_SARS_T_NeoAg_Vax enhanced antigen specific CD8 T cell immunity, which is B cell-dependent. This observation is also supported by previous literature that CD8 T cell immunity is also dependent on B cell and CD4 T cell immunity.^38,45–49,55,73,79,80,91,92^ For instance, One study suggests B/CD4 T cell crosstalk activates Tfh cells, producing IL-21 to improve CD8 responses against cancer.^80^ Since CD4 T cells are also considered to help CD8 T cell responses through various mechanisms^93–98^, B_SARS_T_NeoAg_Vax activated a high level of tumor-specific CD4 T cells, including a large portion of non-follicular CD4 T cells, which may facilitate CD8 T cell response.

Our study also presented a universal nanovaccine platform by incorporating viral B cell epitopes in neoantigen cancer vaccine design, potentially enhancing anticancer efficacy across different types of cancers. The use of viral B cell epitopes offers a safe and universal approach, addressing the challenge of selecting B cell epitopes suitable for various patients with different types of cancers. Furthermore, our study also highlights the critical attributes of nano delivery system to promote B cell-mediated antigen presentation to activate CD4 and CD8 T cell anticancer immunity for enhancing anticancer efficacy. We built two nanovaccine platforms for peptide and mRNA neoantigen vaccine design, where the surface of nanoparticles was conjugated with multivalent SARS-CoV-2 B cell epitopes. The multivalency of viral B cell epitope peptide on the surface of nanoparticles is crucial for BCR crosslinking, antigen uptake, processing, and B cell-mediated antigen presentation to tumor-specific CD4 T cells.^25,58–62^

In summary, our data showed that SARS-CoV-2 B epitope-guided neoantigen cancer nanovaccines enhance anticancer efficacy by promoting antitumor CD4/CD8 T cell antitumor immunity through B cell-mediated antigen presentation. This approach also offers a universal nanovaccine platform for peptide/mRNA neoantigen cancer vaccine design, potentially boosting anticancer efficacy across different types of cancers.

## MATERIALS AND METHODS

### Animal experiments

All animal experiments were conducted according to protocols approved by the University of Michigan Committee on Use and Care of Animals (UCUCA). Animals were maintained under pathogen-free conditions, in temperature- and humidity-controlled housing, with free access to food and water, under a 12-hour light-dark rhythm at the Unit for Laboratory Animal Medicine (ULAM), part of the University of Michigan medical school office of research. Endpoints for anti-tumor efficacy studies were determined using the End-Stage Illness Scoring System, mice receiving an End-Stage Illness Score greater than six were euthanized by CO2 asphyxiation.

### Cells

All cells were maintained at 37 °C in a 5% CO2/95% air atmosphere and approximately 85% relative humidity. Primary B-cells, CD4 T cells and splenocytes were cultured in RPMI-1640 media supplemented with 10% fetal bovine serum, 2-Mercaptoethanol (50 µM) and 1% pen/strep. D2F2/E2 cells generated by cotransfection with pRSV/neo and pCMV/E2 encoding human ErbB-2 (HER2)^99^, were cultured in complete high-glucose DMEM supplemented with 10% NCTC 109 media, 1% L-glutamine, 1% MEM nonessential amino acids, 0.5% sodium pyruvate, 2.5% sodium bicarbonate, 1% pen/strep, 5% cosmic calf serum, and 5% fetal bovine. B16-OVA cells were cultured in RPMI-1640 media supplemented with 10% fetal bovine serum, 2-Mercaptoethanol (54 µM), 1X non-essential amino acids, 10 mM HEPES, 1 mg/ml Geneticin and 1% pen/strep. B16F10 cells were cultured in complete high-glucose DMEM supplemented with 10% fetal bovine serum and 1% pen/strep. KPC 6422 cells obtained from kerafast were cultured in DMEM supplemented with 10% fetal bovine serum, glutamax and 1% pen/strep^100^. Bone marrow-derived dendritic cells (BMDCs) were isolated from mouse bone marrow cells and cultured following previously reported protocols^101,102^.

### Preparation, crosslink and lymph node distribution of Antigen cluster nanovaccine (ACN)

ACN was prepared based on previously reported protocols^65,68^. The core of ACN was made by thermal decomposition and further coated with a polysiloxane-containing copolymer. Au nanodots were then attached onto the surface of ACN core to form an antigen cluster topography. The final Au:Fe ratio of the formulated ACN was quantified by inductively coupled plasma–mass spectrometry (ICP-MS) based on previously reported protocols^103^. Peptides were added to ACN at a 5× weight ratio excess in Milli-Q water and incubated overnight at 4 °C. Peptide loading was determined by fluorescence quantification using a modified fluorescamine peptide quantification assay in the presence of ACN (Ex/Em: 390/465 nm, Biotek Cytation 5)^104^.

To determine the ACN (conjugated to ED-FITC labeled peptide, CDDDPESFDGDPASNTAPLQPEQLQ, 233.6 nmol) distribution, lymph nodes were harvested 12 hours after subcutaneous injections. Lymph nodes were fixed and embedded in optical coherence tomography (OCT) compound and frozen in a CO_2(s)_ + EtOH bath. Tissue sections (15 µm) were prepared, stained and mounted with VECTASHIELD^®^ Mounting Medium for confocal imaging. Brilliant Violet 421 B220, Alexa Fluor^®^ 594 CD 3 and Alexa Fluor^®^ 647 CD169 were used for lymph node immune fluorescence staining.

#### Investigation of BCR crosslink^105^

B cells were isolated from splenocytes of QM mice by EasySep™ mouse B cell isolation kit (STEMCELL). QM B cells (5 * 10^6^ cells/mL) were then incubated with Alexa Fluor 488-affinipure fab fragment goat anti-mouse IgM (20 μg/mL) on ice for 30 minutes in the dark (Jackson: 115-167-020). QM B cells (2 × 10^6^ cells/mL) were then incubated with cy3 labeled and hapten conjugated antigen (20 nM,) in a total volume of 400 µL for 5 minutes at 37 °C. After incubation, cells were immediately fixed with 6% paraformaldehyde (800 µL) for 10 minutes at 37 °C, permeabilized with a 0.1% Triton X HBSS solution (800 µL) for 10 minutes, and then incubated with Alexa Fluor™ Plus 405 phalloidin in staining buffer (200 µL, 5 mg/mL BSA, 0.1% Triton X in HBSS) on ice for 2 hours in the dark. Finally, the cells were washed and plated onto eight-well glass chambers pretreated with 0.1% poly-l-lysine (LabTech II) on ice for at least 4 hours in the dark before confocal imaging.

### Investigation of B/CD4 T cell crosstalk *in vitro*

CD4 T cells were isolated from splenocytes of OT-II mice (The Jackson Laboratory, Strain, #004194) by EasySep™ mouse CD4 T cell isolation kit (STEMCELL). Mice were immunized with SARS-CoV-2 Spike protein (5 µg, Val16-Pro1213, wild type, Genscript) with Alhydrogel® adjuvant 2% (InvivoGen) by intramuscular injection for three times. B cells were isolated from splenocytes and lymph nodes of spike protein immunized mice by EasySep™ mouse B cell isolation kit (STEMCELL). Cells from spleen and lymph node of spike protein immunized mice were depleted B cells by EasySep™ Mouse CD19 positive selection kit II (STEMCELL). CD4, B220, CD69, CD86, CD25 and CFSE are used as markers to measure activation and proliferation. Cell mixtures within each well of 24 well plates including: CD4 T cells (0.5 million, labeled with CFSE, from OT-II mice), B cells (1 million, labeled with CFSE, from spike protein immunized mice), full splenocytes (1.5 million, no CFSE label, from spike protein immunized mice). Cell mixtures are then incubated with different groups for 24h and 96h (All with the same amount of antigens, OT-II CD4 T cell epitope (OVA_323-339_, CISQAVHAAHAEINEAGR, 2 µM) and SARS-CoV-2 B cell epitopes, (CDDDTESNKKFLPFQQFGRDIA, S14P5, 1 µM and CDDDPSKPSKRSFIEDLLFNKV S21P2, 1 µM). After each time point, cells are collected for flow cytometry analysis. For αMHC-II, αICOS and αCD40L antibodies (20 µg/mL) preincubation, all antibodies were preincubated overnight with cells before antigen incubation.

### Analysis of germinal center B cells, antigen-specific germinal center B cells and T follicular helper cells by flow cytometry

Mice were immunized as described in manuscripts. Lymph nodes were harvested for single cell suspension for flow cytometry analysis 10 days after the final immunization. CD3^-^B220**^+^** CD95**^+^** GL-7**^+^** populations were identified as germinal center B cells. Germinal center derived antigen-specific B-cell analysis was measured by tetramer staining based on previously established protocols with minor modifications^106^. Biotin-labeled SARS-CoV-2 peptides were mixed with brilliant violet 421-labeled streptavidin at an 8:1 molar ratio at room temperature for 1 hour to make peptide tetramers. B220^-^CD4^+^ CXCR5^+^ PD-1^+^ populations were identified as Tfh cells. For the αMHC-II (500 µg/mouse), αICOS (200 µg/mouse) or αCD40L (200 µg/mouse) antibodies blocking, the antibodies were intraperitoneal injected 3 days before vaccination, and then dosed 200 µg every two days.

### Analysis of Activation Induced Marker assay for T cells by flow cytometry^70,71^

Mice were immunized as described in manuscripts. Spleen and lymph nodes were harvested for single cell suspension 10 days after the final immunization. Cells (2 million) from different groups were then incubated with B_SARS_-T_HER2_ peptides (2 µg/mL) for 20h at 24 well plates before flow cytometry. B220^-^ CD4^+^ CXCR5^+^ PD-1^+^ populations were identified as Tfh cells. CD69^+^ CD40L^+/-^ populations from Tfh cells were identified as AIM^+^ Tfh cells. B220^-^ CD4^+^ CD69^+^ CD40L^+/-^ populations were identified as AIM^+^ CD4 T cells.

### In vivo anti-tumor efficacy

HER2^+^ breast cancer model: 2.5 × 10^5^ D2F2/E2 cells were subcutaneously inoculated in the right flank of female BALB/c mice at 6 weeks of age. D2F2/E2 cells were prepared at a concentration of 2.5 × 10^6^ cells/mL in 100 µL and were mixed with an equal volume with Matrigel matrix. Pancreatic cancer model: 5 × 10^5^ KPC 6422 cells were subcutaneously inoculated at the right flank of male C57BL/6 mice at 6 weeks of age. Melanoma model: 5 × 10^5^ B16F10 cells were subcutaneously inoculated at the right flank of male C57BL/6 mice at 6 weeks of age. B16-OVA model: 2 × 10^5^ B16-OVA cells were subcutaneously inoculated at the right flank of male C57BL/6 mice or µMt mice at 6 weeks of age. For B cells infusion, 2 × 10^6^ B cells isolated from male C57BL/6 mice at 6 weeks of age by EasySep™ mouse B cell isolation kit (STEMCELL) were infused to male µMt mice 1 day and 3 days before immunization through tail vein injection. For the αMHC-II (500 µg/mouse), αICOS (200 µg/mouse) or αCD40L (200 µg/mouse) antibodies blocking, the antibodies were intraperitoneal injected 3 days before vaccination, and then dosed 200 µg every two days. For the αCD8a (200 µg/mouse) antibody blocking, the antibody was intraperitoneal injected 10 days before vaccination, and then dosed 200 µg every three days. Tumor volumes were calculated as volume = (width)^2^×length/2. End points were determined by using the End-Stage Illness Scoring System.

### Analysis of antigen specific CD4 T and CD8 T cells

2 × 10^6^ cells from lymph nodes or spleen were isolated from each group and incubated with 2 × 10^5^ antigen stimulated BMDC and 2 µg/mL antigen for 20 h and in the presence of Brefeldin A for 4 hours before flow cytometry analysis. Cells were then stained with CD45, CD4 and CD8 markers and permeabilized and fixed by Cyto-Fast™ Fix/Perm Buffer Set (Biolegend). Cells were then stained with IFN-γ antibody and measured by flow cytometry. 5 × 10^5^ splenocytes were isolated from each group and incubated with 2 × 10^5^ antigen stimulated BMDC and 2 µg/mL antigen for 20 h in the ELISPOT plate (R&D systems). ELISPOT measurement and analysis were conducted according to the protocol provided by the manufacturer. Ionomycin and phorbol myristate acetate treated cells were measured as positive control. I-A(b) chicken ova 325-335 QAVHAAHAEIN brilliant violet 421-labeled tetramer, I-A(b) chicken ova 329-337 AAHAEINEA brilliant violet 421-labeled tetramer, I-A(b) chicken ova 328-337 HAAHAEINEA brilliant violet 421-labeled tetramer and H-2K(b) chicken ova 257-264 SIINFEKL PE-labeled tetramer provided by NIH tetramer core facility are used for staining antigen specific CD4 T and CD8 T cells in mice B16-OVA model.

### Single-cell RNA sequencing of tumor samples from pancreatic cancer model

Tumor samples were harvested and dissociated into single-cell suspensions 9 days after the final vaccination. Dead cells were removed using a dead cell removal kit (Miltenyi Biotec). The single-cell suspensions were then stained with TotalSeq™-C Mouse antibody. These suspensions were subjected to final cell counting on a Countess II Automated Cell Counter (Thermo Fisher) and diluted to a concentration of 700-1000 nuclei/µL. We constructed 3′ single-nucleus libraries using the 10x Genomics Chromium Controller and followed the manufacturer’s protocol for 3′ V3.1 chemistry with NextGEM Chip G reagents (10x Genomics). The final library quality was assessed using a TapeStation 4200 (Agilent), and the libraries were quantified by Kapa qPCR (Roche). Pooled libraries were subjected to 150 bp paired-end sequencing according to the manufacturer’s protocol (Illumina NovaSeq 6000). Bcl2fastq2 Conversion Software (Illumina) was used to generate demultiplexed Fastq files, and a CellRanger Pipeline (10x Genomics) was used to align reads and generate count matrices.

### Ionizable lipids synthesis (MMT6-55)

Synthesis of described **Scheme 1**. To a solution of MS-102 (203.0 mg, 0.28 mmol), *N-[(1,1-Dimethylethoxy)carbonyl]-1-methyl-D-tryptophan* (106.8 mg, 0.34 mmol), EDC HCl (64.4 mg, 0.34 mmol) and DMAP (3.5 mg, 0.028 mmol) in dry CH_2_Cl_2_ (5.0 mL), was added at 0°C under Ar. After being stirred at RT for 30 h, the mixture was diluted with CH_2_Cl_2_ and washed with sat. NaHCO_3_ aq and sat. <u>NaCl</u> aq, dried over Na_2_SO_4_, filtered, concentrated, and subjected to silica gel column chromatography (Methanol/Chloromethane= 1/30) to yield **MMT6-52 (88%)**.

^1^H NMR (599 MHz, CDCl_3_) δ 7.56 (d, *J* = 7.9 Hz, 1H), 7.31 – 7.27 (m, 1H), 7.23 (t, *J* = 7.4 Hz, 1H), 7.11 (t, *J* = 7.4 Hz, 1H), 6.90 (s, 1H), 5.11 (d, *J* = 8.3 Hz, 1H), 4.88 (t, *J* = 6.3 Hz, 1H), 4.67 – 4.57 (m, 1H), 4.15 – 4.06 (m, 4H), 3.77 (d, *J* = 2.8 Hz, 3H), 3.50 (d, *J* = 1.6 Hz, 6H), 3.28 (dd, *J* = 8.2, 5.5 Hz, 2H), 2.60 (t, *J* = 6.5 Hz, 2H), 2.40 (q, *J* = 7.0 Hz, 3H), 2.29 (dt, *J* = 9.3, 7.5 Hz, 4H), 1.69 – 1.59 (m, 6H), 1.53 (p, *J* = 6.4, 4.9 Hz, 4H), 1.45 (s, 11H), 1.29 (dd, *J* = 18.4, 7.9 Hz, 46H), 0.90 (td, *J* = 7.0, 2.4 Hz, 8H).

The solution of HCl in dioxane (4 M, 0.2 mL) was added to **MMT6-52** (20 mg, 0.02 mmol) at 0 °C. After stirring for 2 h at RT, the mixture was concentrated to afford **MMT6-55 (99%)**.

^1^H NMR (599 MHz, CDCl_3_) δ 7.60 (d, *J* = 7.8 Hz, 1H), 7.33 (t, *J* = 5.9 Hz, 2H), 7.25 (t, *J* = 7.6 Hz, 1H), 7.14 (t, *J* = 7.4 Hz, 1H), 5.32 (s, 1H), 4.87 (p, *J* = 6.3 Hz, 2H), 4.50 (s, 4H), 4.27 (s, 1H), 4.06 (t, *J* = 6.8 Hz, 3H), 3.82 (s, 3H), 3.70 – 3.56 (m, 3H), 3.51 (s, 1H), 3.39 (s, 2H), 3.26 (s, 2H), 3.11 (s, 8H), 2.32 (dt, *J* = 26.3, 7.2 Hz, 4H), 2.17 – 1.58 (m, 13H), 1.52 (q, *J* = 6.3 Hz, 4H), 1.47 – 1.09 (m, 36H), 0.90 (td, *J* = 7.0, 1.7 Hz, 8H).

### Lipid nanoparticle formulation and characterization

Lipid nanoparticle (LNP) formulation was formulated by mixing an aqueous phase containing the mRNA with an ethanol phase containing the lipids at 3:1 ratio in a microfluidic chip device. The ethanol phase contains a mixture of ionizable lipid (MMT6-55), helper phospholipid (1,2-Distearoyl-sn-glycero-3-PC), cholesterol and PEG-lipid (DMG-PEG2000 or DMG-PEG2000-Maleimide) at predetermined molar ratios. The aqueous phase contains corresponding mRNA (firefly luciferase, enhanced green fluorescent protein, KPC neoantigens) in a 50 mM citrate buffer. The resultant LNP formulations were dialyzed against PBS overnight at 4 °C by Pur-A-Lyzer™ Maxi Dialysis Kit (Millipore Sigma). After dialysis, LNP formulations were concentrated using Amicon ultra-centrifugal filters (MWCO 10KDa, Millipore Sigma) at 5000 rcf to reach the desired volume for experiments. After concentration, SARS-CoV-2 epitopes were incubated with LNP formulations at 2:1 molar ratio between DMG-PEG2000-Maleimide with a gentle stir for 2h at 4°C. The volume-weighted hydrodynamic particle size, and polydispersity index of LNP formulations in Milli-Q water were evaluated with a Malvern Zetasizer Nano-ZS using DLS at 25°C. The encapsulation efficiency of LNP formulations was measured by Quant-iT™ RiboGreen™ RNA Reagent and Kit (invitrogen) followed the protocol provided by the manufacturer.

### mRNA lipid nanoparticle formulations transfection efficiency

Firefly luciferase expression was monitored by incubation of firefly luciferase mRNA encapsulated LNP formulation with DC2.4 cells. DC cells were plated in 24 well plates at 2 × 10^5^ cells/mL overnight and transfected with firefly luciferase mRNA encapsulated LNP formulations (2 ug total mRNA/well) for 24h. Cells were then lysed by Glo lysis buffer and measured by Bright-Glo Luciferase Assay System according to the manufacturer’s protocol (Promega). Enhanced green fluorescent protein (EGFP) expression was monitored by incubation of EGFP mRNA encapsulated LNP formulation with DC2.4 cells. DC cells were plated in the confocal plate at 2.5 × 10^5^ cells/mL overnight and transfected with EGFP mRNA encapsulated LNP formulations (2 ug total mRNA/well) for 6h. Cells were then washed, fixed and imaged by Nikon A1SI confocal.

### In vivo mRNA vaccine anti-tumor efficacy

Mice are immunized with Spike ECD (5 µg, Val16-Pro1213, wild type, Alhydrogel® adjuvant 2%, 1:1 ratio, 100 µL total volume) every 7 days for 3 times to establish pre-existing SARS-CoV-2 epitope immunity. Pancreatic cancer model: 5 × 10^5^ KPC 6422 cells were subcutaneously inoculated at the right flank of male C57BL/6 mice at 6 weeks of age. Mice are immunized with mRNA vaccine (20 µg total mRNA) 4 days after tumor inoculation for 5 times (every 4 days for three doses and every 3 days for the final two doses). Anti-PD-1 are intraperitoneal injected 1 day after vaccination. Tumor volumes were calculated as volume = (width)^2^×length/2. Endpoints were determined by using the End-Stage Illness Scoring System.

### Enzyme-linked immunosorbent assay (ELISA) for antibody titer measurements

Blood was collected 6 days after the final mRNA vaccine immunization. Serum was separated from whole blood by centrifugal separation at 10,000 × g for 5 minutes at 25 °C using Microvette 500 Ser-Gel collection vessels with a clotting activator. Antigen-specific IgG antibody titer was quantified based on previously established protocols for indirect ELISA, with minor modifications^107^. Specifically, SARS-CoV-2 peptides (200 µL, 10 μg/mL in 100 mM carbonate buffer, pH 9.4) were chemically conjugated to ELISA plates through the terminal amine group utilizing Nunc Immobilizer Amino immunoassay plates by overnight incubation with exposure to light at room temperature. Following overnight incubation, ELISA plates were washed three times with 100 mM PBS pH 7.4 with 2% Tween-20. The ELISA plates were then blocked overnight at 4 °C with 300 µL of ELISA blocker (Pierce Protein-Free PBS Blocking Buffer) and washed one time. Serum samples containing primary antibodies were serially diluted (2.5*10^1^ – 5.12*10^4^-fold) using 100 mM PBS pH 7.4 containing 10% ELISA blocker reagent and were added to each well (to 200 µL total volume) for 2 hours incubation at room temperature. After five washes,100 µL of 500-fold diluted anti-IgG-HRP was added to each well and incubated for 1 hour at room temperature. The ELISA plates were washed five times, and then 100 µL of 1-Step Ultra TMB Substrate Solution was added to each well. The solution was allowed to incubate and develop color for 15-20 minutes at room temperature with gentle agitation. Color development was stopped by the addition of 100 µL of 100 mM sulfuric acid. Colorimetric development was quantified by absorbance spectroscopy at 450 nm using a BioTek Cytation 5 plate reader. Antibody titers were determined by any absorbance signal at a given dilution factor that was greater than the PBS control absorbance signal plus three standard deviations.

## Acknowledgments

We thank the use of shared resource facility at the University of Michigan (Pharmacokinetics, Flow cytometry, Microscopy, Advanced Genomics, and In Vivo Animal cores). We thank NIH tetramer core facility for providing tetramers to measure antigen specific CD4/CD8 T cells. We thank M. Cascalho for providing QM mice to measure B cell receptor crosslink. We thank B. Jacobovitz of Microscopy Core for technical assistance. We thank J. Yi for discussion in ordering mRNA sequences. We thank G. Zhu for providing assistance in formulating lipid nanoparticles.

## Author contributions

C.L. designed the project, performed and analyzed all experiments and wrote the paper with all authors; S.M.M.M.T. and M.D. synthesized ionizable lipid for the mRNA studies; Z.M. and A.D. synthesized antigen cluster nanoparticles; F.K. and H.W. helped animal experiments; D.S. and W.G. designed and supervised the project and wrote the paper.

## Data and materials availability

The main data that supports the study are available within the article and supplementary material. The relevant data of this study are available from the corresponding author upon reasonable request. All single cell RNA-sequencing data will be deposited in the GEO database and provided accession number as soon as we have it.

## Reference

1 Ott, P. A. et al. An immunogenic personal neoantigen vaccine for patients with melanoma. Nature 547, 217–221 (2017). 10.1038/nature22991

2 Hu, Z. et al. Personal neoantigen vaccines induce persistent memory T cell responses and epitope spreading in patients with melanoma. Nat Med 27, 515–525 (2021). 10.1038/s41591-020-01206-4

3 Weber, J. S. et al. Individualised neoantigen therapy mRNA-4157 (V940) plus pembrolizumab versus pembrolizumab monotherapy in resected melanoma (KEYNOTE-942): a randomised, phase 2b study. The Lancet 403, 632–644 (2024). 10.1016/S0140-6736(23)02268-7

4 Rojas, L. A. et al. Personalized RNA neoantigen vaccines stimulate T cells in pancreatic cancer. Nature 618, 144–150 (2023). 10.1038/s41586-023-06063-y

5 Hailemichael, Y. et al. Persistent antigen at vaccination sites induces tumor-specific CD8(+) T cell sequestration, dysfunction and deletion. Nat Med 19, 465-+ (2013). 10.1038/nm.3105

6 Liu, S. et al. A DNA nanodevice-based vaccine for cancer immunotherapy. Nature materials 20, 421–430 (2021). 10.1038/s41563-020-0793-6

7 Kissick, H. T. & Sanda, M. G. The role of active vaccination in cancer immunotherapy: lessons from clinical trials. Curr Opin Immunol 35, 15–22 (2015). 10.1016/j.coi.2015.05.004

8 Kissick, H. T. Is It Possible to Develop Cancer Vaccines to Neoantigens, What Are the Major Challenges, and How Can These Be Overcome? Neoantigens as Vaccine Targets for Cancer. Cold Spring Harb Perspect Biol 10 (2018). 10.1101/cshperspect.a033704

9 Finn, O. J. & Rammensee, H. G. Is It Possible to Develop Cancer Vaccines to Neoantigens, What Are the Major Challenges, and How Can These Be Overcome? Neoantigens: Nothing New in Spite of the Name. Cold Spring Harb Perspect Biol 10 (2018). 10.1101/cshperspect.a028829

10 Saxena, M., van der Burg, S. H., Melief, C. J. M. & Bhardwaj, N. Therapeutic cancer vaccines. Nature Reviews Cancer 21, 360–378 (2021). 10.1038/s41568-021-00346-0

11 Hollingsworth, R. E. & Jansen, K. Turning the corner on therapeutic cancer vaccines. NPJ vaccines 4, 7 (2019). 10.1038/s41541-019-0103-y

12 Jiang, T. et al. Tumor neoantigens: from basic research to clinical applications. Journal of hematology & oncology 12, 93 (2019). 10.1186/s13045-019-0787-5

13 Sellars, M. C., Wu, C. J. & Fritsch, E. F. Cancer vaccines: Building a bridge over troubled waters. Cell 185, 2770–2788 (2022). 10.1016/j.cell.2022.06.035

14 Curran, M. A. & Glisson, B. S. New Hope for Therapeutic Cancer Vaccines in the Era of Immune Checkpoint Modulation. Annual review of medicine 70, 409–424 (2019). 10.1146/annurev-med-050217-121900

15 Blass, E. & Ott, P. A. Advances in the development of personalized neoantigen-based therapeutic cancer vaccines. Nature reviews. Clinical oncology 18, 215–229 (2021). 10.1038/s41571-020-00460-2

16 Beura, L. K., Jameson, S. C. & Masopust, D. Is a Human CD8 T-Cell Vaccine Possible, and if So, What Would It Take? CD8 T-Cell Vaccines: To B or Not to B? Cold Spring Harb Perspect Biol 10 (2018). 10.1101/cshperspect.a028910

17 Sahin, U. et al. Personalized RNA mutanome vaccines mobilize poly-specific therapeutic immunity against cancer. Nature 547, 222–226 (2017). 10.1038/nature23003

18 Yarchoan, M. et al. Personalized neoantigen vaccine and pembrolizumab in advanced hepatocellular carcinoma: a phase 1/2 trial. Nat Med 30, 1044–1053 (2024). 10.1038/s41591-024-02894-y

19 Chattopadhyay, S., Chen, J. Y., Chen, H. W. & Hu, C. J. Nanoparticle Vaccines Adopting Virus-like Features for Enhanced Immune Potentiation. Nanotheranostics 1, 244–260 (2017). 10.7150/ntno.19796

20 Bachmann, M. F. & Jennings, G. T. Vaccine delivery: a matter of size, geometry, kinetics and molecular patterns. Nat Rev Immunol 10, 787–796 (2010). 10.1038/nri2868

21 Spohn, G. & Bachmann, M. F. Exploiting viral properties for the rational design of modern vaccines. Expert Rev Vaccines 7, 43–54 (2008). 10.1586/14760584.7.1.43

22 Somiya, M., Liu, Q. & Kuroda, S. Current Progress of Virus-mimicking Nanocarriers for Drug Delivery. Nanotheranostics 1, 415–429 (2017). 10.7150/ntno.21723

23 Bousso, P. T-cell activation by dendritic cells in the lymph node: lessons from the movies. Nature Reviews Immunology 8, 675–684 (2008). 10.1038/nri2379

24 Laumont, C. M., Banville, A. C., Gilardi, M., Hollern, D. P. & Nelson, B. H. Tumour-infiltrating B cells: immunological mechanisms, clinical impact and therapeutic opportunities. Nature Reviews Cancer 22, 414–430 (2022). 10.1038/s41568-022-00466-1

25 Batista, F. D. & Harwood, N. E. The who, how and where of antigen presentation to B cells. Nature Reviews Immunology 9, 15–27 (2009). 10.1038/nri2454

26 Rivera, A., Chen, C. C., Ron, N., Dougherty, J. P. & Ron, Y. Role of B cells as antigen-presenting cells in vivo revisited: antigen-specific B cells are essential for T cell expansion in lymph nodes and for systemic T cell responses to low antigen concentrations. Int Immunol 13, 1583–1593 (2001). 10.1093/intimm/13.12.1583

27 Yuen, G. J., Demissie, E. & Pillai, S. B lymphocytes and cancer: a love-hate relationship. Trends Cancer 2, 747–757 (2016). 10.1016/j.trecan.2016.10.010

28 Qin, Z. et al. B cells inhibit induction of T cell-dependent tumor immunity. Nat Med 4, 627–630 (1998). 10.1038/nm0598-627

29 Shah, S. et al. Increased rejection of primary tumors in mice lacking B cells: inhibition of anti-tumor CTL and TH1 cytokine responses by B cells. Int J Cancer 117, 574–586 (2005). 10.1002/ijc.21177

30 Brodt, P. & Gordon, J. Anti-tumor immunity in B lymphocyte-deprived mice. I. Immunity to a chemically induced tumor. J Immunol 121, 359–362 (1978).

31 Barbera-Guillem, E. et al. B lymphocyte pathology in human colorectal cancer. Experimental and clinical therapeutic effects of partial B cell depletion. Cancer Immunol Immunother 48, 541–549 (2000). 10.1007/pl00006672

32 Shen, P. & Fillatreau, S. Antibody-independent functions of B cells: a focus on cytokines. Nat Rev Immunol 15, 441–451 (2015). 10.1038/nri3857

33 DiLillo, D. J., Yanaba, K. & Tedder, T. F. B cells are required for optimal CD4+ and CD8+ T cell tumor immunity: therapeutic B cell depletion enhances B16 melanoma growth in mice. J Immunol 184, 4006–4016 (2010). 10.4049/jimmunol.0903009

34 de Jonge, K. et al. Inflammatory B cells correlate with failure to checkpoint blockade in melanoma patients. Oncoimmunology 10, 1873585 (2021). 10.1080/2162402X.2021.1873585

35 de Visser, K. E., Korets, L. V. & Coussens, L. M. De novo carcinogenesis promoted by chronic inflammation is B lymphocyte dependent. Cancer Cell 7, 411–423 (2005). 10.1016/j.ccr.2005.04.014

36 Wennhold, K. et al. CD86+ Antigen-Presenting B Cells Are Increased in Cancer, Localize in Tertiary Lymphoid Structures, and Induce Specific T-cell Responses. Cancer Immunology Research 9, 1098–1108 (2021). 10.1158/2326-6066.Cir-20-0949

37 Nielsen, J. S. et al. CD20+ Tumor-Infiltrating Lymphocytes Have an Atypical CD27− Memory Phenotype and Together with CD8+ T Cells Promote Favorable Prognosis in Ovarian Cancer. Clinical Cancer Research 18, 3281–3292 (2012). 10.1158/1078-0432.Ccr-12-0234

38 Cabrita, R. et al. Tertiary lymphoid structures improve immunotherapy and survival in melanoma. Nature 577, 561–565 (2020). 10.1038/s41586-019-1914-8

39 Shimabukuro-Vornhagen, A. et al. Characterization of tumor-associated B-cell subsets in patients with colorectal cancer. Oncotarget 5 (2014).

40 Lu, Y. et al. Complement Signals Determine Opposite Effects of B Cells in Chemotherapy-Induced Immunity. Cell 180, 1081–1097.e1024 (2020). 10.1016/j.cell.2020.02.015

41 Schlößer, H. A. et al. B cells in esophago-gastric adenocarcinoma are highly differentiated, organize in tertiary lymphoid structures and produce tumor-specific antibodies. OncoImmunology 8, e1512458 (2019). 10.1080/2162402X.2018.1512458

42 Lechner, A. et al. Tumor-associated B cells and humoral immune response in head and neck squamous cell carcinoma. OncoImmunology 8, 1535293 (2019). 10.1080/2162402X.2018.1535293

43 Hladikova, K. et al. Tumor-infiltrating B cells affect the progression of oropharyngeal squamous cell carcinoma via cell-to-cell interactions with CD8(+) T cells. J Immunother Cancer 7, 261 (2019). 10.1186/s40425-019-0726-6

44 Yamakoshi, Y. et al. Immunological potential of tertiary lymphoid structures surrounding the primary tumor in gastric cancer. Int J Oncol 57, 171–182 (2020). 10.3892/ijo.2020.5042

45 Helmink, B. A. et al. B cells and tertiary lymphoid structures promote immunotherapy response. Nature 577, 549–555 (2020). 10.1038/s41586-019-1922-8

46 Petitprez, F. et al. B cells are associated with survival and immunotherapy response in sarcoma. Nature 577, 556–560 (2020). 10.1038/s41586-019-1906-8

47 Kroeger, D. R., Milne, K. & Nelson, B. H. Tumor-Infiltrating Plasma Cells Are Associated with Tertiary Lymphoid Structures, Cytolytic T-Cell Responses, and Superior Prognosis in Ovarian Cancer. Clin Cancer Res 22, 3005–3015 (2016). 10.1158/1078-0432.CCR-15-2762

48 Garaud, S. et al. Tumor infiltrating B-cells signal functional humoral immune responses in breast cancer. JCI Insight 5 (2019). 10.1172/jci.insight.129641

49 Hollern, D. P. et al. B Cells and T Follicular Helper Cells Mediate Response to Checkpoint Inhibitors in High Mutation Burden Mouse Models of Breast Cancer. Cell 179, 1191–1206.e1121 (2019). 10.1016/j.cell.2019.10.028

50 Hong, S. et al. B Cells Are the Dominant Antigen-Presenting Cells that Activate Naive CD4(+) T Cells upon Immunization with a Virus-Derived Nanoparticle Antigen. Immunity 49, 695–708.e694 (2018). 10.1016/j.immuni.2018.08.012

51 Liao, W. et al. Characterization of T-Dependent and T-Independent B Cell Responses to a Virus-like Particle. The Journal of Immunology 198, 3846–3856 (2017). 10.4049/jimmunol.1601852

52 Laumont, C. M. & Nelson, B. H. B cells in the tumor microenvironment: Multi-faceted organizers, regulators, and effectors of anti-tumor immunity. Cancer Cell 41, 466–489 (2023). 10.1016/j.ccell.2023.02.017

53 Cattaneo, C. M. et al. Identification of patient-specific CD4+ and CD8+ T cell neoantigens through HLA-unbiased genetic screens. Nature Biotechnology 41, 783–787 (2023). 10.1038/s41587-022-01547-0

54 Katsikis, P. D., Ishii, K. J. & Schliehe, C. Challenges in developing personalized neoantigen cancer vaccines. Nature Reviews Immunology 24, 213–227 (2024). 10.1038/s41577-023-00937-y

55 Chengyi Li, R. C., Luke F. Bugada, Fang Ke, Bing He, Zhixin Yu, Hongwei Chen, Binyamin Jacobovitz, Hongxiang Hu, Polina Chuikov, Brett D. Hill, Syed M. Rizvi, Yudong Song, Kai Sun, Pasieka Axenov, Daniel Huynh, Xinyi Wang, Lana Garmire, Yu Leo Lei, Irina Grigorova, Fei Wen, Marilia Cascalho, Wei Gao, Duxin Sun. Antigen-Clustered Nanovaccine Achieves Long-Term Tumor Remission by Promoting B/CD4 T Cells Crosstalk ACS Nano Accepted (2024).

56 Kaumaya, P. T. B-cell epitope peptide cancer vaccines: a new paradigm for combination immunotherapies with novel checkpoint peptide vaccine. Future Oncol 16, 1767–1791 (2020). 10.2217/fon-2020-0224

57 Wagner, S., Mullins, C. S. & Linnebacher, M. Colorectal cancer vaccines: Tumor-associated antigens vs neoantigens. World J Gastroenterol 24, 5418–5432 (2018). 10.3748/wjg.v24.i48.5418

58 Aikins, M. E., Xu, C. & Moon, J. J. Engineered nanoparticles for cancer vaccination and immunotherapy. Accounts of chemical research 53, 2094–2105 (2020).

59 Cui, C. et al. Neoantigen-driven B cell and CD4 T follicular helper cell collaboration promotes anti-tumor CD8 T cell responses. Cell 184, 6101–6118.e6113 (2021). 10.1016/j.cell.2021.11.007

60 Parker, D. C. T cell-dependent B cell activation. Annu Rev Immunol 11, 331–360 (1993). 10.1146/annurev.iy.11.040193.001555

61 Chen, X. & Jensen, P. E. The role of B lymphocytes as antigen-presenting cells. Arch Immunol Ther Exp (Warsz) 56, 77–83 (2008). 10.1007/s00005-008-0014-5

62 Clark, M. R., Massenburg, D., Siemasko, K., Hou, P. & Zhang, M. B-cell antigen receptor signaling requirements for targeting antigen to the MHC class II presentation pathway. Current Opinion in Immunology 16, 382–387 (2004). 10.1016/j.coi.2004.03.007

63 Poole, A. et al. Therapeutic high affinity T cell receptor targeting a KRAS(G12D) cancer neoantigen. Nat Commun 13, 5333 (2022). 10.1038/s41467-022-32811-1

64 Kreiter, S. et al. Mutant MHC class II epitopes drive therapeutic immune responses to cancer. Nature 520, 692–696 (2015). 10.1038/nature14426

65 Li, C. et al. Antigen-Clustered Nanovaccine Achieves Long-Term Tumor Remission by Promoting B/CD 4 T Cell Crosstalk. ACS Nano 18, 9584–9604 (2024). 10.1021/acsnano.3c13038

66 Amrun, S. N. et al. Linear B-cell epitopes in the spike and nucleocapsid proteins as markers of SARS-CoV-2 exposure and disease severity. EBioMedicine 58 (2020).

67 Tan, Y. S. et al. Mitigating SOX2-potentiated Immune Escape of Head and Neck Squamous Cell Carcinoma with a STING-inducing Nanosatellite Vaccine. Clin Cancer Res 24, 4242–4255 (2018). 10.1158/1078-0432.CCR-17-2807

68 Chen, H. W. et al. Facile Fabrication of Near-Infrared-Resonant and Magnetic Resonance Imaging-Capable Nanomediators for Photothermal Therapy. Acs Appl Mater Inter 7, 12814–12823 (2015). 10.1021/acsami.5b01991

69 Chen, H. W. et al. Highly crystallized iron oxide nanoparticles as effective and biodegradable mediators for photothermal cancer therapy. J Mater Chem B 2, 757–765 (2014).

70 DiPiazza, A. T. et al. COVID-19 vaccine mRNA-1273 elicits a protective immune profile in mice that is not associated with vaccine-enhanced disease upon SARS-CoV-2 challenge. Immunity 54, 1869–1882.e1866 (2021). 10.1016/j.immuni.2021.06.018

71 Reiss, S. et al. Comparative analysis of activation induced marker (AIM) assays for sensitive identification of antigen-specific CD4 T cells. PloS one 12, e0186998 (2017). 10.1371/journal.pone.0186998

72 Meylan, M. et al. Tertiary lymphoid structures generate and propagate anti-tumor antibody-producing plasma cells in renal cell cancer. Immunity 55, 527–541.e525 (2022). 10.1016/j.immuni.2022.02.001

73 Overacre-Delgoffe, A. E. et al. Microbiota-specific T follicular helper cells drive tertiary lymphoid structures and anti-tumor immunity against colorectal cancer. Immunity 54, 2812–2824.e2814 (2021). 10.1016/j.immuni.2021.11.003

74 Fridman, W. H. et al. B cells and tertiary lymphoid structures as determinants of tumour immune contexture and clinical outcome. Nature Reviews Clinical Oncology 19, 441–457 (2022). 10.1038/s41571-022-00619-z

75 Fridman, W. H., Sibéril, S., Pupier, G., Soussan, S. & Sautès-Fridman, C. Activation of B cells in Tertiary Lymphoid Structures in cancer: Anti-tumor or anti-self? Seminars in Immunology 65, 101703 (2023). 10.1016/j.smim.2022.101703

76 Oliveira, G. et al. Phenotype, specificity and avidity of antitumour CD8(+) T cells in melanoma. Nature 596, 119–125 (2021). 10.1038/s41586-021-03704-y

77 Hu, Z. et al. Personal neoantigen vaccines induce persistent memory T cell responses and epitope spreading in patients with melanoma. Nat Med 27, 515–525 (2021). 10.1038/s41591-020-01206-4

78 Bruno, T. C. et al. Antigen-Presenting Intratumoral B Cells Affect CD4+ TIL Phenotypes in Non–Small Cell Lung Cancer Patients. Cancer Immunology Research 5, 898–907 (2017). 10.1158/2326-6066.Cir-17-0075

79 Sagiv-Barfi, I., Czerwinski, D. K., Shree, T., Lohmeyer, J. J. K. & Levy, R. Intratumoral immunotherapy relies on B and T cell collaboration. Sci Immunol 7, eabn5859 (2022). 10.1126/sciimmunol.abn5859

80 Cui, C. et al. Neoantigen-driven B cell and CD4 T follicular helper cell collaboration promotes anti-tumor CD8 T cell responses. Cell 184, 6101–6118 e6113 (2021). 10.1016/j.cell.2021.11.007

81 Gabrilovich, D. Mechanisms and functional significance of tumour-induced dendritic-cell defects. Nat Rev Immunol 4, 941–952 (2004). 10.1038/nri1498

82 Sica, A. & Bronte, V. Altered macrophage differentiation and immune dysfunction in tumor development. J Clin Invest 117, 1155–1166 (2007). 10.1172/JCI31422

83 Italiano, A. et al. Pembrolizumab in soft-tissue sarcomas with tertiary lymphoid structures: a phase 2 PEMBROSARC trial cohort. Nat Med 28, 1199–1206 (2022). 10.1038/s41591-022-01821-3

84 Biswas, S. et al. IgA transcytosis and antigen recognition govern ovarian cancer immunity. Nature 591, 464–470 (2021). 10.1038/s41586-020-03144-0

85 Mazor, R. D. et al. Tumor-reactive antibodies evolve from non-binding and autoreactive precursors. Cell 185, 1208–1222.e1221 (2022). 10.1016/j.cell.2022.02.012

86 Mandal, G. et al. IgA-Dominated Humoral Immune Responses Govern Patients’ Outcome in Endometrial Cancer. Cancer Research 82, 859–871 (2022). 10.1158/0008-5472.Can-21-2376

87 Bod, L. et al. B-cell-specific checkpoint molecules that regulate anti-tumour immunity. Nature 619, 348–356 (2023). 10.1038/s41586-023-06231-0

88 Oh, D. Y. et al. Intratumoral CD4(+) T Cells Mediate Anti-tumor Cytotoxicity in Human Bladder Cancer. Cell 181, 1612–1625 e1613 (2020). 10.1016/j.cell.2020.05.017

89 Corthay, A. et al. Primary antitumor immune response mediated by CD4+ T cells. Immunity 22, 371–383 (2005). 10.1016/j.immuni.2005.02.003

90 Ferris, S. T. et al. cDC1 prime and are licensed by CD4(+) T cells to induce anti-tumour immunity. Nature 584, 624–629 (2020). 10.1038/s41586-020-2611-3

91 Wieland, A. et al. Defining HPV-specific B cell responses in patients with head and neck cancer. Nature (2020). 10.1038/s41586-020-2931-3

92 Chaurio, R. A. et al. TGF-β-mediated silencing of genomic organizer SATB1 promotes Tfh cell differentiation and formation of intra-tumoral tertiary lymphoid structures. Immunity 55, 115–128.e119 (2022). 10.1016/j.immuni.2021.12.007

93 Bennett, S. R., Carbone, F. R., Karamalis, F., Miller, J. F. & Heath, W. R. Induction of a CD8+ cytotoxic T lymphocyte response by cross-priming requires cognate CD4+ T cell help. J Exp Med 186, 65–70 (1997). 10.1084/jem.186.1.65

94 Bennett, S. R. M. et al. Help for cytotoxic-T-cell responses is mediated by CD40 signalling. Nature 393, 478–480 (1998). 10.1038/30996

95 Schoenberger, S. P., Toes, R. E. M., van der Voort, E. I. H., Offringa, R. & Melief, C. J. M. T-cell help for cytotoxic T lymphocytes is mediated by CD40–CD40L interactions. Nature 393, 480–483 (1998). 10.1038/31002

96 Bourgeois, C., Rocha, B. & Tanchot, C. A Role for CD40 Expression on CD8^+^ T Cells in the Generation of CD8^+^ T Cell Memory. *Science (New York*, N.Y.) 297, 2060–2063 (2002). doi:10.1126/science.1072615

97 Janssen, E. M. et al. CD4+ T-cell help controls CD8+ T-cell memory via TRAIL-mediated activation-induced cell death. Nature 434, 88–93 (2005). 10.1038/nature03337

98 Ferris, S. T. et al. cDC1 prime and are licensed by CD4+ T cells to induce anti-tumour immunity. Nature 584, 624–629 (2020). 10.1038/s41586-020-2611-3

99 Whittington, P. J. et al. Her-2 DNA versus cell vaccine: immunogenicity and anti-tumor activity. Cancer Immunol Immunother 58, 759–767 (2009). 10.1007/s00262-008-0599-x

100 Li, J. et al. Tumor Cell-Intrinsic Factors Underlie Heterogeneity of Immune Cell Infiltration and Response to Immunotherapy. Immunity 49, 178–193.e177 (2018). 10.1016/j.immuni.2018.06.006

101 Lutz, M. B. et al. An advanced culture method for generating large quantities of highly pure dendritic cells from mouse bone marrow. Journal of immunological methods 223, 77–92 (1999). 10.1016/s0022-1759(98)00204-x

102 Song, Y. et al. Albumin nanoparticle containing a PI3Kγ inhibitor and paclitaxel in combination with α-PD1 induces tumor remission of breast cancer in mice. Sci Transl Med 14, eabl3649 (2022). 10.1126/scitranslmed.abl3649

103 Clauson, R. M., Chen, M., Scheetz, L. M., Berg, B. & Chertok, B. Size-Controlled Iron Oxide Nanoplatforms with Lipidoid-Stabilized Shells for Efficient Magnetic Resonance Imaging-Trackable Lymph Node Targeting and High-Capacity Biomolecule Display. ACS Appl Mater Interfaces 10, 20281–20295 (2018). 10.1021/acsami.8b02830

104 Udenfriend, S. et al. Fluorescamine: a reagent for assay of amino acids, peptides, proteins, and primary amines in the picomole range. *Science (New York*, N.Y.) 178, 871–872 (1972). 10.1126/science.178.4063.871

105 Veneziano, R. et al. Role of nanoscale antigen organization on B-cell activation probed using DNA origami. Nature nanotechnology 15, 716–723 (2020). 10.1038/s41565-020-0719-0

106 Franz, B., May, K. F., Jr., Dranoff, G. & Wucherpfennig, K. Ex vivo characterization and isolation of rare memory B cells with antigen tetramers. Blood 118, 348–357 (2011). 10.1182/blood-2011-03-341917

107 Sanchez, A. B. et al. A general process for the development of peptide-based immunoassays for monoclonal antibodies. Cancer chemotherapy and pharmacology 66, 919–925 (2010). 10.1007/s00280-009-1240-1

